# CoSTA: Unsupervised Convolutional Neural Network Learning for Spatial Transcriptomics Analysis

**DOI:** 10.1101/2021.01.12.426400

**Authors:** Yang Xu, Rachel Patton McCord

**Author notes:** Correspondence (R.P.M.).

## Abstract

The rise of spatial transcriptomics technologies is leading to new insights about how gene regulation happens in a spatial context. Here, we present CoSTA: a novel approach to learn spatial similarities between gene expression matrices via convolutional neural network (ConvNet) clustering. By analyzing simulated and previously published spatial transcriptomics data, we demonstrate that CoSTA learns spatial relationships between genes in a way that emphasizes whole patterns rather than pixel-level correlation. CoSTA provides a quantitative measure of how similar each pair of genes are by their spatial pattern rather than only classifying genes into categories. We find that CoSTA identifies narrower, but biologically relevant, sets of significantly related genes as compared to other approaches.

## Background

Spatial transcriptomics has recently gained extensive attention from the scientific community. Different technologies have enabled high resolution measurements of how gene regulation is spatially organized across a tissue or thousands of single cells.[1] However, current analysis pipelines often treat each pixel in an expression matrix as an independent feature, thus losing spatial information. For example, the seqFISH+ technique can fluorescently detect 10,000 mRNAs in situ at single cell resolution, and there are often groups of cells that have correlated gene expression with their neighbors to make up larger structures. However, the original report analyzed these expression patterns using PCA and hierarchical clustering, treating each cell as an independent feature, rather than preserving spatial positions of cell neighbors.[2] Slide-seq similarly produces high-throughput spatially resolved transcription information, using sequencing rather than fluorescence. Previous analyses of Slide-seq data first identified spatially non-random gene expression, but then looked for genes expressed in similar patterns using pixel-level overlap analysis rather than according to spatial features.[3] Existing algorithms for analysis of spatial transcriptomics are based on statistical modeling and primarily propose to distinguish spatially expressing or variable (SE or SV) genes from random spatial expression noise. For example, both SpatialDE and SPARK analysis approaches estimate how significant the spatial pattern of a gene is.[4, 5] SpatialDE further builds in an unsupervised pattern detection algorithm to cluster significant SE genes into different groups which have certain spatial patterns in collective. SPARK, in contrast, was designed only for finding SE genes. To examine spatial relationships between genes, this method still relies on hierarchical clustering that uses individual pixels as features. Therefore, even though SPARK can identify genes with significant spatial patterns, the latter part of the SPARK analysis decouples the expression from its original spatial context. Thus far, existing spatial transcriptomics analyses involve either multi-step complex feature engineering for spatial quantification or human-imposed rigid or statistical modeling-based screening of candidate SE genes. In the existing methods, the similarity of expression pattern between two genes is either binary--whether or not the genes cluster together--or is quantified based on pixel-level correlation.

In this work, we propose an approach inspired by computer vision and image classification to find relationships between spatial expression patterns of different genes while preserving the full spatial context (Fig. 1a). Our goal is to find quantitative comparisons between gene expression patterns in a way that preserves spatial relationships between neighboring cells and tissue regions. We aim for a method that will recognize an overall similar shape of expression, even if certain sets of pixels are not exactly overlapping. This is conceptually similar to image recognition in computer vision tasks. The use of convolutional neural networks brought a success of deep learning in computer vision and have demonstrated a wide range of applications, including image classification and object recognition. A few groups have proposed different approaches to use convolutional neural networks (ConvNet) in unsupervised learning.[6-8] Thus, here, we adopt an unsupervised **Co**nvNet learning strategy for **S**patial **T**ranscriptomics **A**nalysis (CoSTA). With simulated data, we show that CoSTA can correctly classify a variety of different spatial patterns and that the patterns CoSTA is detecting depend on spatial groupings rather than individual pixels. Then, we apply CoSTA to published MERFISH and Slide-seq data and show that CoSTA sometimes identifies smaller sets of genes with significant spatial relationships, but these identified relationships are biologically relevant.

**Fig 1.**
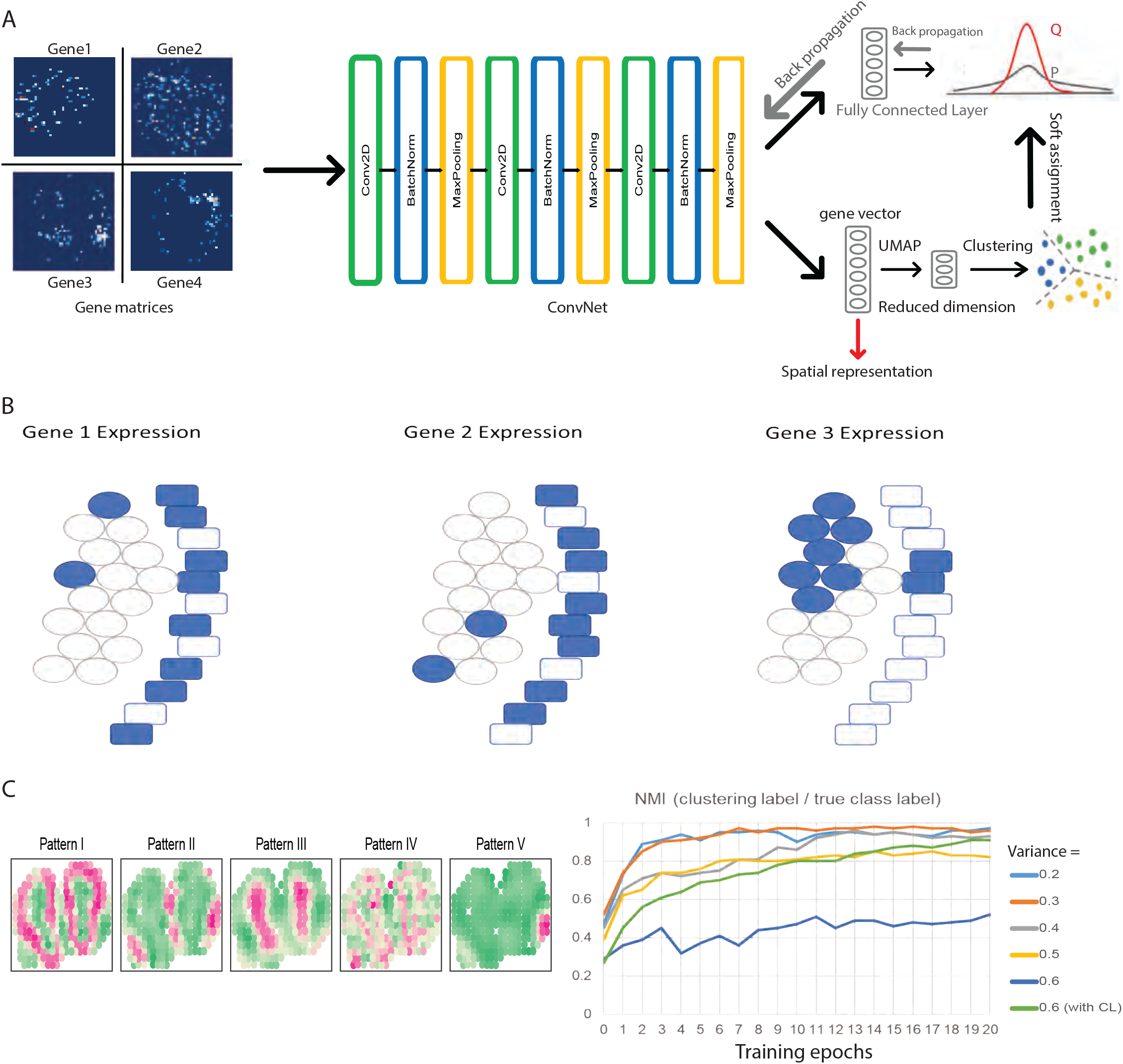
CoSTA model approach and motivation. a, Overall CoSTA pipeline. Inputs are gene matrices from spatial transcriptomic experiments. ConvNet stage forwards images through 3 convolutional layers and then flattens the output into a spatial representation vector. UMAP reduces dimensionality of the spatial representations from the ConvNet stage before these gene representations are used to cluster genes with GMM. Each gene is then assigned cluster probabilities based on distances to cluster centroids, which are transformed to an auxiliary target distribution that can be minimized by reducing bi-tempered logistic loss and/or center loss. Gradients are backpropagated through a fully connected layer to ConvNet. The process is repeated until the model converges, at which point the output from the ConvNet is used as the final spatial representation (red arrow). b, Biologically-inspired example in which overlap does not capture all aspects of spatial pattern similarity. Rectangles represent an epithelial cell layer while ovals represent stromal cells. By overlap comparison, Gene 1 has the same similarity to both Gene 2 and Gene 3 (40% overlap). However, the biologically relevant expression along the epithelial layer is only shared between Gene 1 and Gene 2. Detecting this shape similarity requires learning a spatial representation. c, Performance of CoSTA in synthetic datasets. left panel: 5 real expression patterns in mouse olfactory bulb data replicate 11; right panel: learning curves of CoSTA classifying simulated genes belonging to these 5 patterns with different noise levels. NMIs are measured between clustering labels by CoSTA and the true class label.

## Results

### CoSTA architecture: training a ConvNet with pseudo-labels generated by GMM clustering

Though there are many unsupervised learning strategies, we chose to apply the workflow of DeepCluster, because it is straightforward and easy to implement.[6] Our CoSTA approach consists of two main parts: clustering by Gaussian mixture model (GMM) and weight updating as commonly performed in training neural networks (Fig. 1a and see Methods for detailed description). Our inputs are sets of gene expression images, where each image is the matrix recording the expression levels of one gene at each position in space and all images belong to the same biological space. We first initialize a ConvNet randomly and then forward these gene expression matrices through the ConvNet. Our ConvNet consists of three convolutional layers, and each convolutional layer is followed by a batch normalization layer and a max pooling layer. We flatten the matrix output from the last max pooling layer into a vector that captures the spatial features of the gene expression data. The size of this vector will vary depending on the image size from a given spatial transcriptomics technique. We then apply L2-normalization across features and reduce dimensionality by UMAP before we perform GMM clustering of genes. UMAP can preserve global and local structures during dimension reduction and previously showed better performance in image clustering than other dimension-reduction methods such as Isomap and t-SNE.[7, 9] The purpose of this clustering is to generate labels so that we can update the ConvNet as in other common supervised neural network training approaches. When the ConvNet is randomly initialized, features extracted by this ConvNet are weak. However, using them to generate labels can still guide the ConvNet to learn more discriminative features. Indeed, Caron et al. showed DeepCluster can learn from weak signal to bootstrap the discriminative power of a ConvNet.[6] Instead of giving each gene a single cluster label, we assign an auxiliary target distribution as a soft assignment. This approach emphasizes genes with high confidence in the clustering task and discounts noisy labels persisting from the random initialization of ConvNet. Doing this can also lead to more stable target values for training the neural network.[8] Finally, we use these soft assignments to train the ConvNet. We add a fully connected layer after the ConvNet that produces probabilities for each gene being assigned to each label. Thus, we can optimize the model by minimizing bi-tempered logistic loss based on Bregman Divergences between the soft assignments from GMM clustering and the probabilities from the fully connected layer.[10] In summary, the CoSTA approach uses a ConvNet clustering architecture which repeats 1) generating features by ConvNet, 2) generating soft assignments by GMM clustering, and 3) using soft assignments to update ConvNet. Once we finish training, we only retain the trained ConvNet for the purpose of feature extraction. Since the ConvNet mainly consists of convolutional layers, the final vector for each gene extracted by ConvNet should be a spatial representation. Using this spatial representation, we can then quantify the relationship between any two genes within one spatial transcriptomics dataset, visualize all SE genes in this dataset by UMAP, and assign patterns through common clustering algorithms. Further details about the rationale of this learning architecture can be found in Methods.

### Rationale for using spatial patterns rather than exact pixel overlap

To demonstrate the spatial information lost by overlap analysis and why a spatial representation approach such as CoSTA is useful, we present a simplified biologically-inspired conceptual example (Fig. 1b). In biological tissue sections, we commonly observe structures such as a tightly connected epithelial layer of cells (rectangles in the cartoon) adjacent to a collection of stromal cells (circles). In this example, the spatial expression patterns of three genes are shown. Comparing gene expression patterns by overlap only, we observe that Gene 1 and 2 have the same amount of overlap as Gene 1 and 3 (40%). Thus, an overlap approach to measure gene pattern similarity, like the one used in previous Slide-seq analysis, would report that Gene 1 is equally similar to both Gene 2 and Gene 3.[3] However, biologically, it is relevant that Gene 1 and Gene 2 are expressed primarily in the epithelial layer while Gene 3 is expressed in the stroma. This biological difference is not detected by strict overlap, but instead requires a spatial representation that would detect the vertical stripe of epithelial layer expression as a salient pattern. In computer vision, filters are commonly used to find this kind of local correlation, and the success of ConvNet in pattern recognition also relies in the use of filters for identifying local correlations. Using signals of how these 3 genes respond to the same filters, a ConvNet approach will identify Gene 1 and 2 as more similar and Gene 3 as less similar. Therefore, we are motivated to use our ConvNet clustering based CoSTA approach to prioritize similar shape more than overlap for biological cases where layers of cells and the overall patterns of groups of cells matter more than independent individual cell identities.[11]

### Tests on synthetic data show CoSTA’s high specificity, reliance on spatial relationships, and ability to distinguish signal from noise

As a first test of CoSTA’s ability to detect correlated spatial patterns in the absence of exact overlap, we use the MNIST handwritten digit image data.[12] When the aim is to find which digits have correlated handwritten patterns to the digit 3, CoSTA identifies only other instances of digit 3 as correlated (100% specificity). In contrast, overlap analysis finds some samples of all other digits as correlated digits of 3 (58% specificity) (Fig. S1). Meanwhile, CoSTA identifies a smaller subset of the digit 3s as correlated (35% sensitivity) while overlap analysis captures more correlated digits overall (65% sensitivity) in its less specific set (Fig. S1). As shown below, this increased specificity but possibly decreased sensitivity of CoSTA compared to other techniques appears to hold true in biological data as well.

Before applying CoSTA to real spatial transcriptomics data, we next tested its performance on 5 synthetic datasets, simulated based on real expression patterns from mouse olfactory bulb, following the simulation method in SPARK (Fig. 1c left panel)[5, 13]. We generated 2,000 fake gene expression matrices for each pattern, to mimic data for 10,000 total genes. To simulate noise and variability for each gene, we added residual errors onto each spatial coordinate independently based on a normal distribution with mean of zero and variance ranging from 0.2 to 0.6. We then evaluated whether CoSTA could assign each simulated noisy gene to the correct pattern. When CoSTA was initialized, the Normalized Mutual Information (NMI) against the true class label ranged from 0.27 to 0.57 (Fig. 1c right panel). As training proceeded, CoSTA learned discriminative features to distinguish the 5 patterns, eventually achieving NMIs from 0.85 to 0.98 against the true class label (Fig. 1c right panel, Table S1). For the highest noise level (0.6) we found that combining both center loss (CL) and bi-tempered logistic loss during CoSTA training substantially improved CoSTA’s accuracy (NMI increased from 0.52 to 0.91). However, CL pushes samples to those centroids and is only applicable when the final number of patterns is known. Thus, we do not include CL for biological situations.

To demonstrate that CoSTA learns spatial rather than pixel-level patterns from these synthetic datasets, we shuffled the pixel positions in these synthetic datasets. Shuffling all the gene matrices exactly the same way keeps the pixelwise overlap information identical while disrupting correlations between neighboring pixels, thus destroying the spatial pattern. If a pattern detection method is successfully using spatial relationships between neighboring pixels, its ability to classify patterns should be disrupted by this kind of shuffling. Indeed, we found that CoSTA cannot distinguish the genes into correct pattern labels as well with shuffled data (NMI ranges from 0.32 to 0.89), demonstrating that CoSTA is detecting spatial features that depend on the positions of neighboring pixels, rather than features that can be captured by a set of single pixels (Fig. S2 and Table S1). When we applied the CoSTA model trained at 0.4 noise level to progressively more shuffled images, we found that the ability to classify genes into groups declined proportional to the amount of shuffling (Figure S3A). We also tested SpatialDE on these true and shuffled synthetic datasets. SpatialDE performed very well on the true datasets, as expected. However, shuffling the data did not usually change the performance of SpatialDE (Table S1), indicating an important difference between CoSTA and SpatialDE: SpatialDE is more likely to detect patterns of individual pixels while CoSTA emphasizes the spatial positions of these pixels relative to each other and overall shapes of patterns.

Using this same synthetic data, we next performed a disruption test to demonstrate a disadvantage of using individual pixels as features to analyze spatial transcriptomics data. For half of the simulated gene matrices, we masked a certain region of the pattern, and the masked region doesn’t change expression pattern visually (Fig. S3b). This mimics a situation in which a certain region is obscured or not sampled well for technical reasons experimentally. Using pixel overlap to identify patterns, in this case, assigns masked and unmasked genes into separate groups, even though they otherwise belong to the same pattern. In contrast, CoSTA is resistant to this disruption (Fig. S3b).

In real spatial transcriptomics data, not all genes will belong to a clear spatial pattern— some genes that are not relevant to the given tissue or condition may only yield random noise or be fairly uniformly expressed. To mimic this situation, we further followed the simulation approach in SPARK to generate synthetic datasets that have 5 spatial patterns and have mixed SE (spatially expressed) and non-SE genes (Fig. S4). We trained CoSTA on these data with different ratios of SE and non-SE genes, from 90:10 to 10:90. We found that the representation of SE genes by CoSTA is distinct from non-SE genes, even when CoSTA was trained with a high percentage of non-SE genes. Meanwhile, CoSTA demonstrates the capacity to distinguish different patterns of SE genes even when non-SE genes exist (Fig. S4). Further, CoSTA does not separate even a large number of non-SE genes into separate categories, showing that it does not create false signal out of noise. Here, we also note that a strength of CoSTA compared to methods like SpatialDE is that the output feature vector enables visualization, as is presented throughout these simulation results. While Spatial DE can classify genes into categories, it does not produce a result that can visualize how SE and non-SE genes are separated as we did here for CoSTA. Overall, the performance of CoSTA with synthetic data demonstrates that CoSTA can learn discriminating spatial features.

### CoSTA classifies genes by cell type and identifies quantitative relationships between genes in MERFISH data

To extend the application of CoSTA to real spatial transcriptomics data, we first applied it to reanalyze a MERFISH dataset (see MERFISH Analysis in Methods for complete details).[14] In order to compare with published analyses using the SPARK approach, we focused on the same slice of the mouse hypothalamus (Bregma + 0.11 mm from animal 18).[5] The expression patterns of a set of 155 genes expected to be spatially variable were measured with MERFISH for this slice, along with 5 blank control genes. We first initialized a ConvNet and forwarded the MERFISH spatial gene expression matrices through it to obtain gene feature vectors. Then we clustered the 155 spatially variable genes with the 5 blank genes and with 9 cell type-specific expression patterns defined by the original publication through a combination of MERFISH and scRNA-seq data. We clustered these genes, controls, and cell type patterns into 10 groups and visualized them by UMAP. Without training, SE genes, control genes, and cell types are spread across the 2-dimensional UMAP space and boundaries between groups are not distinctively defined (Fig. 2a). Next, we trained the CoSTA model to obtain refined feature vectors. After training, SE genes, control genes and cell types formed distinct groups that have clearer boundaries in the 2D visualization (Fig. 2b) and refined cluster memberships that reproducibly and quantitatively form tighter clusters according to a linear intrinsic dimensionality (LID) estimator (Fig. 2c) [15].

**Fig 2.**
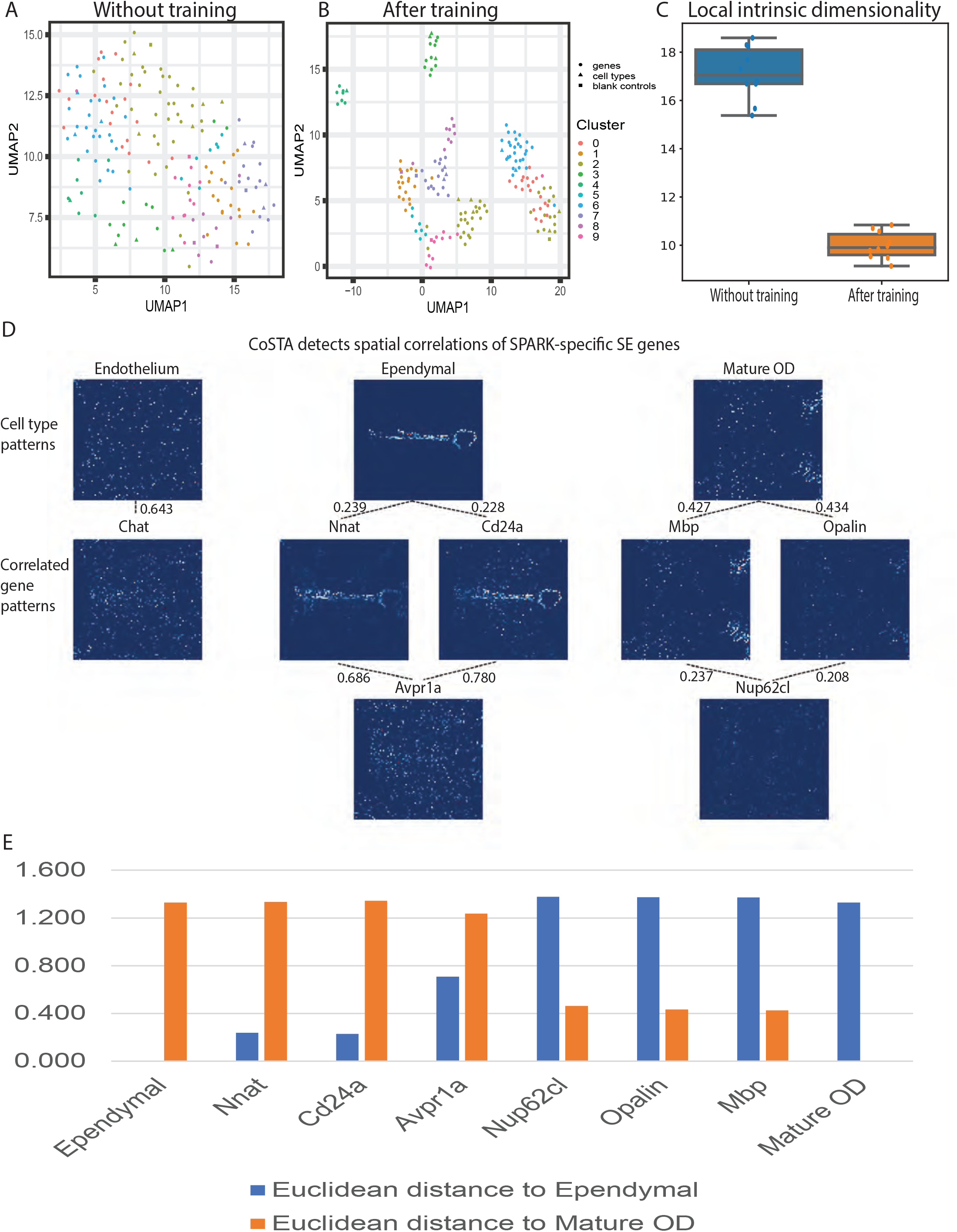
Analyzing MERFISH data with the CoSTA approach. a and b, Visualization of the spatial feature vectors obtained for each gene, blank controls, and reference cell type patterns from MERFISH data in a 2D UMAP layout. Filled circles represent true genes, filled triangles represent cell-types, and filled squares are blank controls. a, features extracted from a randomly initialized ConvNet with no training. Each dot is a gene, blank control, or cell type pattern. Colors indicate cluster labels obtained from clustering on the full CoSTA-derived feature vectors; b, features extracted by trained ConvNet. Each dot is colored with the original clustering labels from *a* to show how some cluster memberships rearrange. c, Local intrinsic dimensionalities of spatial representations by CoSTA without and after training (10 independent runs of CoSTA). d, CoSTA-detected spatial correlations of genes identified as SE uniquely by SPARK. Top row displays known cell type specific expression patterns for 3 cell types. Lower rows display expression patterns of particular genes. Dotted lines indicate CoSTA-identified similarity between a pair of genes or a gene with a cell type pattern. Raw count values for each image are scaled from 0 to 1 to normalize the visual comparison. e, Euclidean distances between each gene shown in d and Ependymal or Mature OD patterns. Euclidean distances are measured using the CoSTA spatial representation.

From this MERFISH data, SPARK identified 145 SE genes including one blank control, and SpatialDE found 139 SE genes with one blank control.[5] CoSTA is designed primarily to detect similarities between spatial gene expression patterns, rather than to estimate spatial relevance (identify SE genes). So, to define which genes are called SE by CoSTA, we examined which genes CoSTA identified as highly correlated to one of the 9 pre-defined cell type specific expression patterns. We found a correlation threshold at which CoSTA identified 133 SE genes associated with one of the different cell type patterns, while none of the blank controls were called associated with a pattern (Table S2). Thus, CoSTA’s sensitivity is slightly lower than SPARK and SpatialDE, but with higher specificity (no blank controls detected). However, CoSTA’s result is both more sensitive and more specific than the Trensceek approach, which only identified 108 SE genes and one blank control.[16]

Three genes in this MERFISH dataset, *Avpr1a, Chat*, and *Nup62cl*, were highlighted by Sun et al., because they were only identified as SE by SPARK.[5] CoSTA is able to identify the spatial expression patterns of these genes, but also reveals by quantitative similarity that these genes are more distantly related to cell type expression patterns than other genes. We examined both the significantly similar groups determined by CoSTA and used the spatial representation learned by CoSTA to measure Euclidean distances of these genes to each other and to cell type expression patterns (Fig. 2D, E and Table S2). For example, CoSTA identifies genes such as *Nnat* and *Cd24a* as significantly similar to the Ependymal cell type pattern (Dotted lines, Fig. 2D). *Avpr1a* is quantified as more distant from this Ependymal pattern (Fig. 2E), though it does show some similarity to *Nnat* and *Cd24a* (Fig. 2D). Similarly, *Mbp* and *Opalin* are significantly correlated to the Mature OD cell type pattern (Fig. 2D, E and Table S2). *Nup62cl* is more distant from the Mature OD than *Opalin* and *Mbp*, but is related to the expression patterns of *Mbp* and *Opalin*. Visual inspection of *Avpr1a* and *Nup62cl* confirms that these patterns are quite noisy and less similar to the key cell type pattern (Fig. 2d). Thus, by quantifying relationships between patterns rather than reporting uniform sets of SE genes, CoSTA clarifies that these genes are likely identified by SPARK and no other method because they are in fact less spatially similar to key cell type patterns. CoSTA’s ability to quantify relationships between genes, rather than only categorizing genes, is important in biological situations, where there is often going to be a range of relative similarity that would be oversimplified by strict categorization.

### CoSTA learns spatial pattern-dependent representations of Slide-seq data

We next expand our application of CoSTA to Slide-seq data. Slide-seq takes advantage of high-throughput single cell RNA sequencing and barcoding. Therefore, it enables spatial gene expression measurement for all genes in the genome.[3] As a first demonstration that CoSTA can be applied to this type of high-throughput spatial transcriptomics data, we performed an experiment-mixing test to evaluate whether CoSTA can separate different spatial patterns. Due to the unavailability of a “gold standard” for positive and negative spatial similarity of gene expression, we mixed gene matrices from four different spatial transcriptomics experiments by Slide-seq and tested the ability of CoSTA to deconvolve them.[3] Each overall experiment is performed on an independent brain slice of a different mouse, so the shapes and spatial features of each experimental sample overall constitute a large difference between experiments. Each gene within each experiment will have a somewhat different pattern (and it will be our next goal to distinguish those differences and similarities), but we first tested whether genes within the same experiment could be classified together based on their overall spatial features. We implemented training as above and then clustered the mixed experiment gene matrices into 4 clusters. The confusion matrix shows clustering labels are largely consistent with true experimental labels (Table S3).

We next performed a shuffling test on gene matrices from one Slide-seq experiment, to break correlated patterns of neighboring regions in the way described for the shuffling of synthetic data above. We trained a new model and examined model-reported similarity among expression patterns of ten random genes. If CoSTA successfully learned spatial features that distinguish the expression of these genes, the distances between two genes should change when spatial

patterns and relationships between neighboring pixels are disrupted. We randomly selected *Prdx5* as the reference gene and calculated Euclidean distances of 9 other genes with it. We order these ten genes based on their distances to *Prdx5*. Then, we shuffled gene matrices 100 times, passed the shuffled matrices through the trained ConvNet, and recalculate paired distances with *Prdx5* (Fig.3a). We find that in 5 of 9 comparisons, distances decreased upon shuffling, as the distinctive patterns captured by CoSTA were removed by shuffling, converting the matrices into generic, more similar patterns. In 4 of 9 comparisons, distances increased with shuffling, likely indicating that key similarities between the spatial patterns became disrupted during shuffling (Fig. 3b). In contrast, the similarity measured by overlap analysis would not change after shuffling since individual pixels were shuffled identically. This result again suggests, this time using real biological data, that the learned features by CoSTA are strongly tied to the spatial expression pattern.

**Fig 3.**
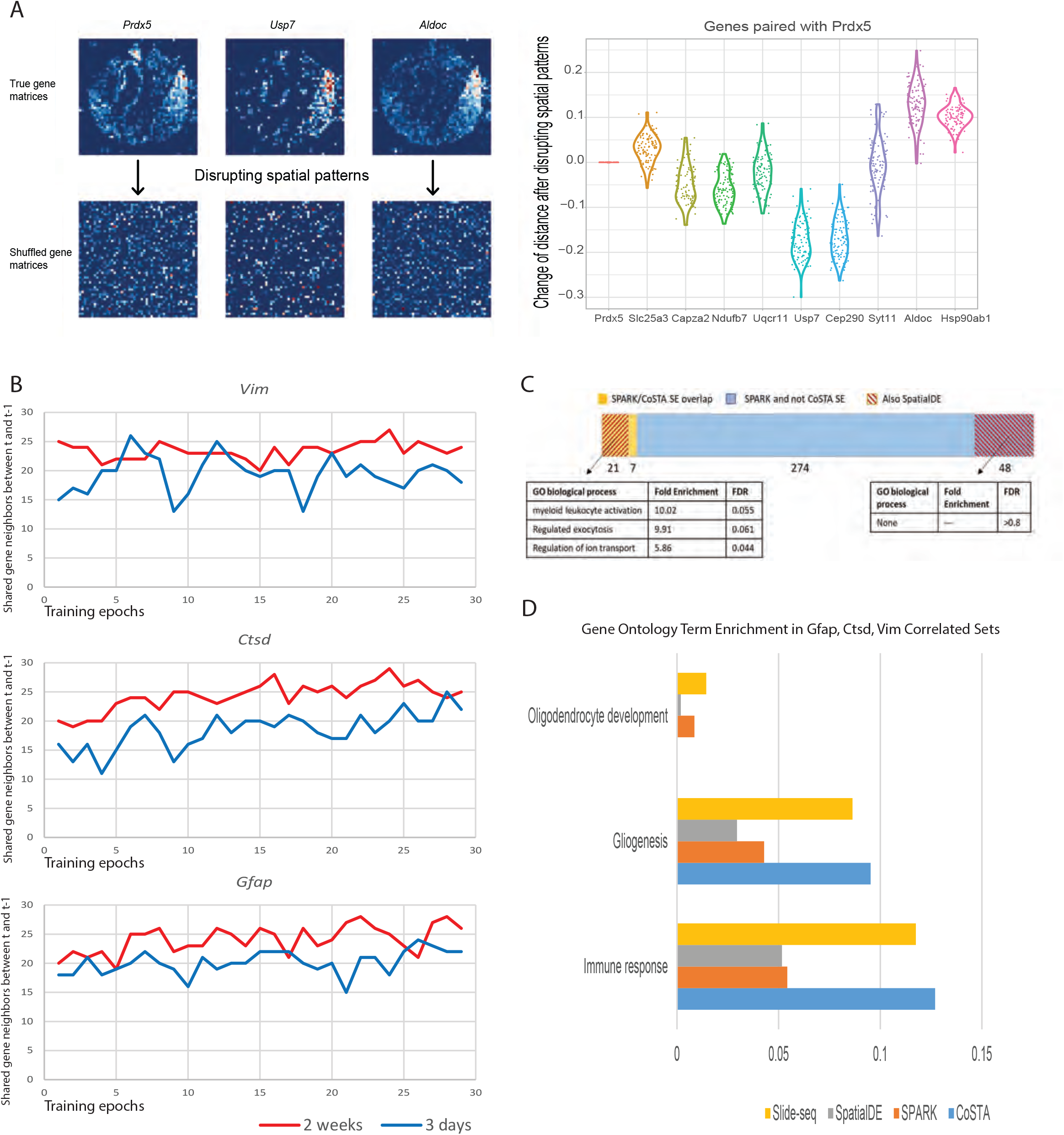
CoSTA Analysis of Slide-seq data. a, Shuffling test to disrupt spatial patterns. Left panel: The first row shows the three original spatial expression patterns of three example genes. Images in the second row are spatial patterns after shuffling (all images shuffled in the same way so that pixel-level overlap is preserved while spatial neighbor relationships are broken). Right panel: Distances between 9 randomly selected genes and *Prdx5*. Genes are ordered based on how close they are to *Prdx5* using spatial features extracted by CoSTA from true gene matrices (left to right: closest to farthest). Shuffled gene matrices are forwarded through CoSTA, and distances between gene pairs are subtracted from the unshuffled distances. Each point represents distance change for one shuffling (100 shufflings total). Red line at 0 indicates no change in distance would be observed using overlap calculations. b, The number of overlapped gene neighbors of *Vim, Ctsd*, and *Gfap* before and after each weight updating across all training epochs (30 nearest neighbors considered, see Fig. S5 for different size neighbor sets). Results shown for two experiments: 3 days (blue) or 2 weeks (red) after brain injury. c, Proportions of SPARK and SpatialDE-identified genes that overlap with CoSTA SE genes. Yellow = SPARK correlated genes also SE by CoSTA. Blue = SPARK correlated genes not SE by CoSTA. Red cross-hatching = Proportion of each category also identified as correlated by SpatialDE. Below, Gene Ontology enrichment (Panther) of genes that overlap between SPARK, SpatialDE AND CoSTA (left) and those that overlap between SPARK and Spatial DE but NOT CoSTA (Right). d, GO term enrichment in the correlated gene sets from different approaches for biologically relevant functions identified by the original Slide-seq analysis. Quantified along the axis is the fraction of genes in each method’s correlated gene list that are annotated with the given GO term.

### Ensemble learning identifies stable relationships between spatial gene expression patterns

We next applied CoSTA to reanalyze two spatial transcriptomics datasets measured by Slide-seq.[3] These datasets are derived from two biological conditions: 3 days after brain injury (“3 days”) and 2 weeks after brain injury (“2 weeks”). In the first investigation of these two datasets in Slide-seq, Rodriques et al. primarily focused on genes that were spatially correlated with *Vim, Ctsd* and *Gfap* at both 3 days and 2 weeks after brain injury.[3] For comparison, we also examined genes correlated with *Vim, Ctsd* and *Gfap* from our CoSTA results. One property of our approach is that features of each gene change every epoch when weights are updated. This may result in changes to the nearest neighbors of a gene during model training and can be used to infer how strong and stable the inferred spatial relationships are in a given condition. We measured the overlap between detected *Vim, Ctsd, and Gfap* neighbor genes before and after weight updating across training epochs, and we found neighbors tend to be more stable for the 2 weeks dataset than for the 3 days dataset (Fig. 3b and Fig. S5). This may indicate that in the acute phase after injury, genes related to *Vim, Ctsd* and *Gfap* are more variable and less spatially patterned, but these patterns become stronger at 2-week time point after injury.

To screen significantly spatially patterned genes out from noise, we use ensemble learning. Briefly, we initialized 5 ConvNets and trained them separately. We then calculated the nearest neighbors for every gene in the same dataset, at neighbor set sizes of 5, 10, 15, 20, 25, 30, 40, 50, and 100. We use a broad range of neighboring levels because different genes may form different sizes of communities. Next, we calculated Jaccard similarities across the 5 CoSTA models and keep genes that have an averaged Jaccard similarity larger than 0.2 at least in one level. We call genes that pass the threshold “stable”, and genes that are filtered out as “unstable”. We propose that the percentage of stable vs. unstable genes represents the degree of spatial patterning in the experiment set. Overall, a smaller proportion of genes were considered stable at 3 days, consistent with the more variable gene neighbors observed for the 3-day condition above. These ‘stable’ genes can also be considered a CoSTA-derived set of ‘spatially expressed’ (SE) genes for comparison to SE genes identified by SPARK. The majority of CoSTA-SE genes are also called SE by SPARK (86% at 3 days and 78% at 2 weeks, Fig. S6a). *Vim, Ctsd*, and *Gfap* are considered SE by CoSTA in the 2-week data but not in the 3-day dataset. Notably, *Vim, Ctsd*, and *Gfap* are also not present in the 3 days SE gene list identified by SPARK, and only *Ctsd* and *Gfap* were identified as SE genes by SPARK in the 2 weeks data. We note that less strongly patterned genes could reflect actively variable biological regulation (such as might happen during acute response to injury), not only technique noise. We are unable to definitively distinguish a weak spatial pattern from inherent noise, because of lack of “ground truth” for pattern matching. However, we can, as above, disrupt spatial patterns by shuffling the true datasets, maintaining pixelwise correlations between genes but removing spatial information. We shuffled a whole set of gene matrices from 3 days and 2 weeks and applied CoSTA to these datasets. As when we shuffled simulated data in Figure S2, we find that this shuffled dataset has overall lower NMI than its original dataset during training (Fig. S6b; see Methods for details of NMI use). Further, substantially fewer SE genes are identified in the 2 week randomized data as compared to the real data (Fig. S6c). This again demonstrates that CoSTA captures spatial features that are distinct from individual pixel information. For true 3-day and shuffled 3-day data, there is not a clear difference in the number of identified SE genes (Fig. S6c). This again suggests that the spatial patterns are much less strong within the 3-day dataset. Indeed, few patterns are visually obvious for example gene matrices from 3 days (Fig. S7a).

### CoSTA identifies smaller, but specific and biologically relevant, sets of spatially correlated genes compared to SPARK and SpatialDE

We focused our further analysis on the 2-week data. We applied SpatialDE and SPARK to this dataset for comparison to CoSTA. The original Slide-seq publication previously identified 843 genes that are correlated with *Vim, Ctsd*, and *Gfap* via overlap analysis.[3] However, CoSTA, with a rigid neighbor similarity stability threshold, identified many fewer correlated genes (63 with z-scores < -2.325), and only 19 genes matched the original Slide-seq set (Fig. S8a). SPARK first identified 1294 significantly SE genes and then clustered them into 10 groups by hierarchical clustering with individual pixels as features. Our CoSTA correlated gene list only has 5 gene overlaps with genes that are grouped with *Vim, Ctsd*, and *Gfap* by SPARK. We also used SpatialDE to find significant SE genes. Surprisingly, the whole dataset passed the SpatialDE test for significant spatial expression. Then, we applied the unsupervised pattern detection algorithm built in SpatialDE to cluster genes into 10 groups. This resulted in a large number of genes grouped with *Vim, Ctsd*, and *Gfap*. A majority of our CoSTA set (41 genes) overlaps with genes identified by SpatialDE (Fig. S8a). The set of correlated genes identified by CoSTA is much smaller than sets identified by the other 3 methods. This is in part because CoSTA requires stable relationships between neighboring genes to be classified as an SE gene at all, and only SE genes can then be identified as highly similar to the genes of interest. Indeed, out of 350 genes identified by SPARK as correlated with *Vim, Ctsd, and Gfap*, only 28 are even classified as SE genes by CoSTA. However, we observe evidence that this CoSTA-identified SE subset is reliable and meaningful. First, 75% of these CoSTA-SE genes identified by SPARK are also identified as correlated by SpatialDE, while in the remaining non CoSTA-SE set, there is only a 15% overlap between SPARK and SpatialDE (Fig. 3c). Further, the genes overlapping between SPARK, SpatialDE, and CoSTA have biologically relevant function enrichment (such as ion transport and exocytosis) while the genes overlapping between SPARK and SpatialDE but not identified as SE by CoSTA show no function enrichment at all (Fig. 3c). We also observe visible evidence of spatial pattern similarity to the 3 genes of interest among genes considered SE and highly correlated by CoSTA and less evidence of similarity for genes identified only by SPARK (Fig. S8b).

Further, we find that the 63 genes identified by CoSTA as significantly correlated to *Vim, Gfap*, and *Ctsd* are highly enriched for meaningful biological function. In the original study, Rodriques et al. highlighted that genes correlated with *Vim, Ctsd*, and *Gfap* are enriched for functions in immune response, gliogenesis and oligodendrocyte development—all functions that are biologically expected in response to injury.[3] We found that the correlated genes identified by CoSTA have higher enrichment in immune response and gliogenesis than the genes identified by SpatialDE, SPARK and this original Slide-seq report (Fig. 3d). However, none of genes fall into category of oligodendrocyte development. When we visually inspected expression patterns of genes in the category of oligodendrocyte development, their individual and collective patterns do not have similarities to expression patterns of *Vim, Ctsd, and Gfap*. They are either noisy or expressed globally (Fig. S7b). From results above, we conclude that CoSTA returns a reduced, stringent set of correlated genes that are more enriched for biological significance than the larger sets returned by other methods.

As noted earlier, one key difference between CoSTA and these other methods is that CoSTA provides not only sets of similar genes, but also quantitative pairwise comparisons between all genes. Thus, we can extract from CoSTA a ranked list of how similar each CoSTA-SE gene is to *Vim, Ctsd*, and *Gfap* (Table S4). This enables us to search for enriched biological functions using similarity rankings rather than an arbitrary cutoff using the GOrilla enrichment tool.[17] Using the whole ranked list, we find novel enriched functions such as collagen metabolism, astrocyte differentiation, and vascular endothelial growth factor signaling that may be relevant to damage repair (Fig. S8c). Pixelwise correlation can also be used to create a ranked similarity list. When these two approaches are compared, we observe some highly ranked genes shared by both approaches with clear pattern overlap to the query genes. Where the two approaches differ widely in gene ranking, CoSTA-specific genes tend to have the key patterns of expression as contiguous patterns superimposed on a generic weak background while pixel-specific genes tend to have isolated pixels overlapping the key areas (Fig. S8d).

To avoid observation bias by only looking at a few example genes that are classified differently by different approaches, we next globally compared the types of spatial patterns detected uniquely by CoSTA and other previous methods. For each method (CoSTA, SpatialDE, SPARK, and the original Slide-seq overlap approach), we consider the list of genes classified as spatially correlated with *Vim, Ctsd, and Gfap* as described above. The average expression pattern of genes detected as correlated to these query genes varies somewhat according to approach. Notably, the average CoSTA pattern is more localized to the upper right region, where the damage was induced (Fig. 4a). In contrast, SPARK, SpatialDE, and Slide-seq each identify so many correlated genes that their average pattern looks very much like the average pattern of all genes in the dataset (compare Fig. 4a with Fig. S9) rather than distinctive. This again emphasizes the smaller, but perhaps more specific, set of genes identified as correlated by CoSTA. When we compare genes identified as correlated by CoSTA and not certain other techniques, we can see that CoSTA-unique genes have certain local patterns that were not captured as much by other methods (Fig. 4b). In contrast, again, genes detected by other methods and not CoSTA look more similar on average to the average gene expression of the entire gene set (Fig. 4c).

**Fig 4.**
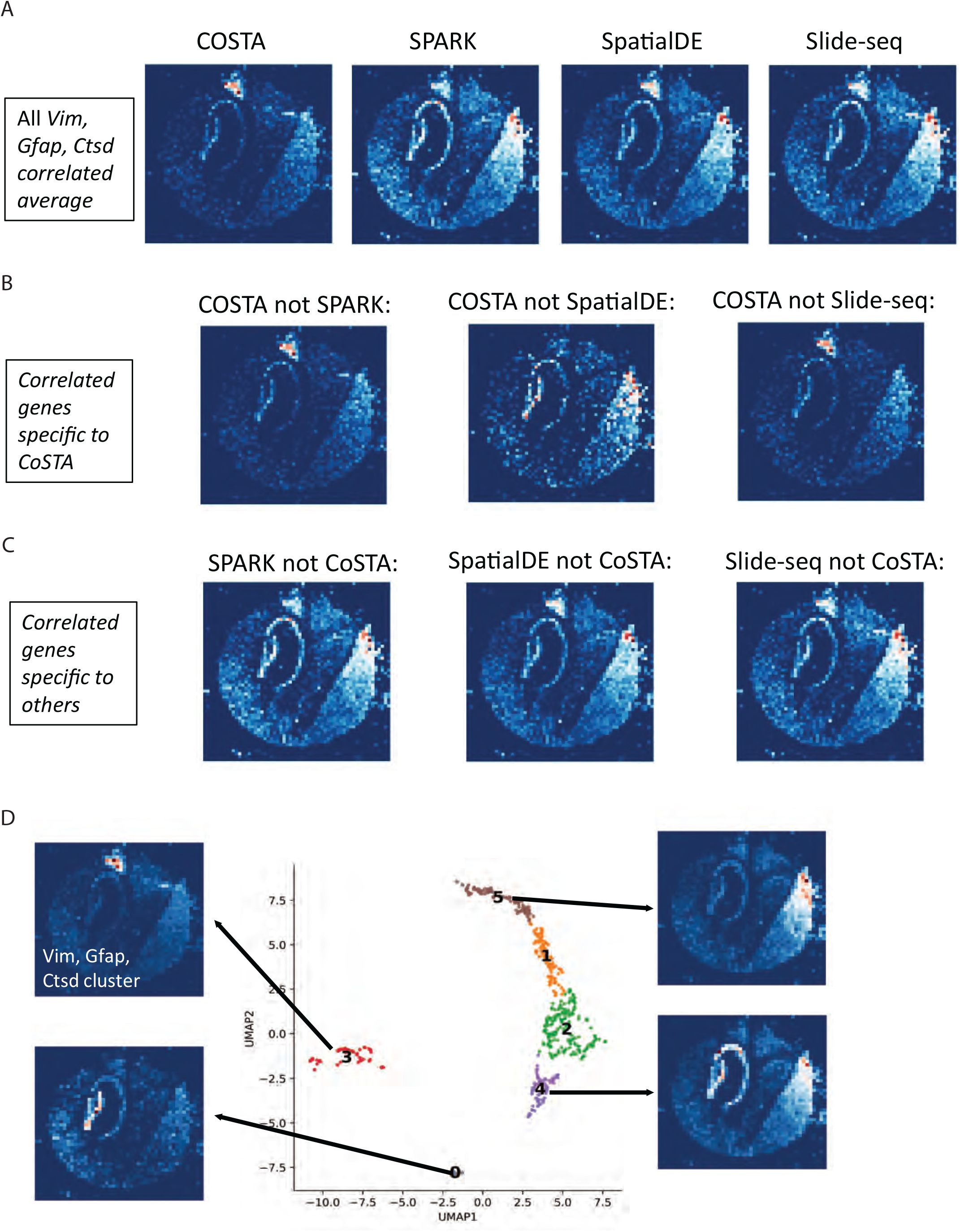
Collective expression patterns detected by different methods in Slide-seq data. a, Average expression patterns of *Vim, Gfap, Ctsd* and their correlated genes after 2 weeks brain injury defined by different methods. b, average expression patterns of *Vim, Gfap, Ctsd* correlated genes detected by CoSTA and not by the indicated methods. c, Average expression patterns of *Vim, Gfap, Ctsd* correlated genes detected by other methods and not CoSTA. d, Gene clusters of CoSTA SE genes at 2-week time point on 2 UMAP dimensions. Mean expression patterns are presented for selected clusters. Visualization with raw count values that are scaled from 0 to 1.

Finally, rather than using a significant correlation threshold, we clustered all CoSTA-determined SE genes at the 2-week time point into 6 groups using the learned spatial representation. The cluster that contains *Vim, Ctsd*, and *Gfap* (cluster 3) is composed of 89 genes expressed in a distinct pattern (Fig. 4d and Table S5). Other clusters also successfully identify distinctive spatial patterns of expression (Fig. 4d and Fig. S9). We also used SpatialDE to cluster SE genes identified by CoSTA into 6 clusters. We found that the two methods share many commonalities in detecting patterns, with some disagreements (Fig. S9). Notably, when only the narrower set of SE genes identified by CoSTA is used, the cluster of genes identified by SpatialDE containing *Vim, Gfap*, and *Ctsd* (cluster 2, Fig. S9) has a much more specific, localized pattern than when SpatialDE default settings are used to classify all genes. This again suggests that CoSTA provides a meaningful increase in specificity by identifying genes with stable spatial relationships.

## Discussion

We have shown that our CoSTA approach can successfully implement deep learning ideas from computer vision to infer spatial gene expression relationships. Identifying spatial patterns from high-throughput spatial transcriptomics data is still challenging, however. We often do not have a clear ground truth answer for what should be detected as a pattern vs. noise and what similarities in patterns are most biologically relevant. Different approaches will have different strengths and weaknesses depending on the types of patterns and relationships to be detected. The very first step in any approach to analyzing spatial transcriptomics data is estimating significant SE genes. To identify SE genes, SpatialDE relies on the assumption that spatial expression of a given gene follows a multivariate normal distribution across spatial coordinates.[4] However, this assumption leads all genes in a Slide-seq dataset to be identified as SE genes by SpatialDE. This may occur because noisy signals generated by the Slide-seq experiment may also follow or are confounded within the multivariate normal distribution. Therefore, a multivariate normal model will not be able to distinguish spatial patterns from noise in certain types of experimental data. Different from SpatialDE, both SPARK and CoSTA make use of kernels to identify SE genes. SPARK defined 5 periodic and 5 gaussian kernels to cover a range of possible spatial patterns that the authors believe are observed in common biological datasets.[5] Therefore, identifying SE genes involves a statistical evaluation of how well kernels match spatial patterns of interest. This SPARK approach is very valuable if an experimental dataset is accompanied by prior knowledge about relevant spatial patterns. Kernels in CoSTA also serve a similar purpose but are not predefined. Instead, kernels in CoSTA are learned through training a neural network. To identify SE genes, we rely on the idea that a true spatial pattern should be collective, which means a group of genes should share a spatial pattern. Therefore, when we apply kernels learned independently from 5 ConvNets, genes in the same group should have similar responses to these kernels. Conversely, a noisy gene expression pattern would respond to the 5 sets of ConvNet kernels differently, clustered with different groups of genes each time. Indeed, we showed that this kernel approach helps identify a more focused set SE genes in Slide-seq data without requiring an *a priori* definition of relevant patterns that SPARK requires. We have shown by various measures that the SE genes identified by CoSTA are a much smaller set, but with high enrichment for meaningful biological function, and more likely to be also detected by multiple other methods, increasing confidence in this set.

Identification of SE genes is just the beginning of extracting biological meaning from spatial gene expression. Careful analysis of the spatial relationships between genes is also necessary. Often, as in overlap analysis, studying gene relationships is based on vectorizing gene expression patterns and measuring their similarities in a latent space without considering spatial information such as the position of neighboring datapoints. One key motivation for CoSTA, therefore, is to preserve a spatial and shape representation of gene expression patterns. In comparison, SPARK does not have a pattern detection function, but can be combined with hierarchical clustering with pixels as features to assign each gene a pattern label. SpatialDE implements a clustering model based on a spatial Gaussian-process-based (GP) prior.[4] This clustering model is an extension of GMM with the addition of a spatial prior on cluster centroids. Therefore, pattern detection by SpatialDE goes beyond the pixel level. In our method, we define the key goal as learning a spatial representation for each gene. We have demonstrated that features learned by CoSTA are not isolated to individual pixels, while SpatialDE responds more to individual pixel information in our simulations. Because of use of convolutional layers, spatial features learned by our method represent local patterns and multiple local patterns together form the global pattern for the gene matrix. Finally, vectorizing gene matrices allows us not only to find different spatial patterns within a dataset by clustering but also to study spatial relationships of pairs of genes. Such a pairwise examination, in contrast, is not implemented in SpatialDE.

Not only in detection of a narrower set of SE genes, but also in identifying relationships between genes, our results consistently suggest that CoSTA provides more specific though less sensitive results than other methods. Throughout our analyses, we find that overlap approaches, as well as SPARK and SpatialDE tend to group together larger sets of genes that are more distant in their spatial pattern relationships, while CoSTA captures a narrower and more specific set of genes. This was observed in our analysis of digit image data as well as in applications to Slide-Seq and, to a lesser extent, MERFISH. This difference in outcomes again demonstrates the different advantages and disadvantages of different approaches. CoSTA would likely be more useful in a case where users want to narrow their set of candidate related genes for future experiments. We also note throughout the Methods section alterations to parameters of CoSTA that could allow for detection of more general patterns.

Again, depending on the biological reality underlying the data, different approaches will have different advantages. The CoSTA approach will have advantages in cases where overall pattern shape is important, while direct overlap calculations may perform better when exact cell to cell correlation is more biologically relevant. The CoSTA approach may also have future applications to datasets in which images of different genes are not from the identical biological section, but instead from neighboring tissue slices, as is common in traditional histology. If a pattern or shape of expression is maintained while exact overlap is lost, as we demonstrated with our simulated masking approach, CoSTA could still detect such a pattern similarity where an overlap approach would not.

## Conclusions

In this study, we demonstrated that our deep learning CoSTA approach provides a different angle to spatial transcriptomics analysis by focusing on the shape of expression patterns. CoSTA includes more information about the positions of neighboring pixels than does an overlap or individual pixel correlation approach. CoSTA can be applied to any form of spatial transcriptomics data that are represented in matrix form to find genes expressed in similar patterns as well as to evaluate the strength of the spatial patterning of each gene. We find that CoSTA captures more focused groups of spatially related genes while still detecting the biological function information found by other approaches that report larger sets of related genes.

## Methods

### Resize Gene Images and Normalization

The raw images of Slide-seq consist of over 1,000,000 pixels, which makes computation difficult. Therefore, we first binned 100 pixels into one pixel and resized matrices from different experiments into the same 48×48 image size. This results in a lower resolution, which may obscure small-scale fine details, but large scale global features of expression patterns of genes are preserved. CoSTA can be applied to any spatial transcriptomics dataset at any resolution, as long as the user has sufficient computational resources available. To avoid extreme computational burden, we recommend that users interested in high resolution features zoom into regions of interest and crop images in that region to efficiently apply CoSTA to their data. After binning, we normalized gene matrices as described in Svensson et al.[4] This normalization involves finding the total gene expression counts for each pixel across all gene matrices and then normalizing each pixel of each matrix by the log total counts across all matrices for this pixel. If this normalization is not performed, the expression of a gene could be over or undercounted at certain spatial locations where expression levels were systematically high or low for all genes. Normalization by total counts at each pixel ensures that our approach captures the spatial covariance for each gene beyond this potential artifactual effect. For visualization of expression patterns, we instead use averaged raw count values, and scale values from 0 to 1 divided by the maximum value. Thus, expression images in all figures are in 0 to 1 scale. This allows a more direct visual inspection of the raw data.

### CoSTA Architecture

#### 1. ConvNet

The ConvNet stage of CoSTA consists of 3 convolutional layers for Slide-seq and MERFISH analysis. Inputs are sets of spatial gene expression images (matrices) as described above. We first initialize a ConvNet randomly and then forward these gene expression matrices through the ConvNet. All weights in convolutional layers are initialized on a Xavier uniform distribution. Each convolutional layer is activated by a rectified linear unit function and is followed by a batch normalization layer and a max pooling layer to reduce the size of the output. To produce a feature vector for each gene, we flatten the matrix output from the last max pooling layer by concatenating all matrix columns into a single column. One fully connected layer is added to the model after the last max pooling layer with a customized softmax activation to produce outputs as probabilities (See **4. Loss Function**). The fully connected layer is only used during training, when we need gradients to pass backwards through the model. Once trained, this fully connected layer will be discarded, and we use L2-normalized outputs as the spatial representations. Specific parameters used in ConvNet, such as the number and size of filters in each convolutional layer, can be found in python code. We note that different numbers of convolutional layers have been used for different image classification tasks. We recommend that users start with a 3-convolutional-layer network for initial data exploration. However, if a dataset has a larger size of gene matrices, outputs from the 3-convolutional-layer network will be very long vectors. Therefore, users can increase the number of convolutional layers to decrease the dimensions of outputs if needed.

#### 2. UMAP and Clustering

The flattened spatial representation vector output from the three convolutional layers is reduced by UMAP before GMM clustering. We implemented UMAP using the original python source code[9]. We set up “n_neighbors=20” and “min_dist=0”, while using UMAP for dimension reduction. To cluster samples into N clusters, a user can reduce dimensions to N UMAP-dimensions. In this study, we reduce all samples to 30 UMAP-dimensions and cluster all samples into 30 clusters by GMM. While 30 clusters are used here for the model training purpose, once the model is trained, the user can use the final output vector of spatial features to cluster genes into any number of groups desired. To test the influence of the initial choice of number of clusters, we tested 10, 20, and 30, 50, 75, and 100 clusters in 2-week Slide-seq data. Using larger numbers of clusters leads to the identification of fewer SE genes (Fig. S10a). Our model can converge no matter how many clusters are used for training (Fig. S10b). For a purpose of comparison, we called the 15 nearest genes of *Vim, Ctsd*, and *Gfap* individually, and total 45 genes in one test as correlated genes were used for comparing effects of the number of clusters. The choice of the number of clusters will influence the scale of correlated expression pattern detected (Fig. S10c). More global pattern differences will be detected using smaller numbers of clusters while finer scale pattern distinctions are detected with larger numbers of clusters (Fig. S10c). Increasing the number of clusters will also bring a disadvantage of larger computational cost and longer training time (Table S6). In this case, 30 clusters show good specificity, and the detected spatial pattern is not further refined with increasing cluster numbers (Fig. S10c). Without ground truth for a dataset, the number of clusters must be chosen based on the scale of patterns desired to be detected for a particular biological application and the results inspected visually.

#### 3. Auxiliary Target Distribution as Soft Assignment

After clustering, we calculate centroids by averaging samples in the same cluster (Eq. 1).

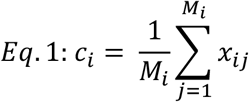

Where *c*_*i*_ is the centroid for the *i*^*th*^cluster, *M*_*i*_ is the total number of samples in this cluster, and *x*_*i,j*_is a reduced UMAP vector for the *j*^*th*^sample in the *i*^*th*^cluster.

Then, each sample is assigned probabilities based on Euclidean distances to cluster centroids (Eq. 2).

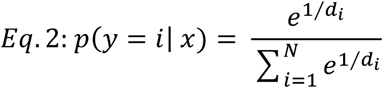

Where *d*_*i*_ is the Euclidean distance of sample *x* to the centroid *c*_*i*_, and *N* is the total number of clusters.

Next, we transform probabilities of each sample to an auxiliary target distribution using equation 3.

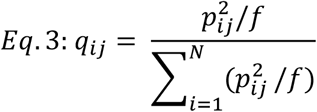

where 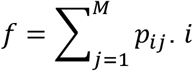 denotes the *i*^*th*^cluster and *j*denotes the *j*^*th*^sample, *p*_*ij*_is probability that the *j*^*th*^sample belongs to the *i*^*th*^that we get through Equation 2. *q*_*ij*_is the auxiliary target probability that the *j*^*th*^sample belongs to the *i*^*th*^cluster. This transformation was proposed by Xie et al, which is raising *p*_*ij*_to the second power and then normalizing by frequency per cluster.[18] The use of power 2 is to highlight samples that have high confidence in the clustering task and discount samples for which the model is uncertain about cluster assignment.

#### 4. Loss Function

To optimize the neural network, we use bi-tempered logistic loss based on Bregman Divergences as the primary loss function. Bi-tempered logistic loss was proposed by Amid et al and showed advantage of making supervised learning robust to noise.[10] To achieve the robustness, they devised tempered softmax function and tempered logistic loss and gave detailed mathematical reasons behind (Eq. 4 and 5). We reason that training CoSTA also faces the problem of unknown noise within the data, because clustering will assign wrong labels to samples. This is even true when clustering is based on the ConvNet that is randomly initialized. Therefore, use of bi-tempered logistic loss is to deal with wrong or uncertain labels generated by clustering. When both *t*_1_and *t*_2_are equal to 1, bi-tempered logistic loss is the common KL-divergence loss with softmax activation.

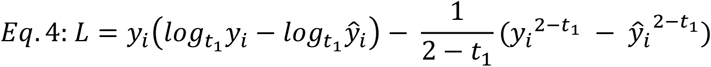

Where 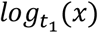 can approximate to 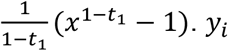 is the target value and ŷ_i_ is the predicted value out of the fully connected layer.

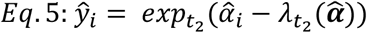

Where 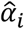 is linear activation of output of the fully connected layer for the *i*^*th*^cluster, and 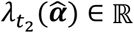 is s.t.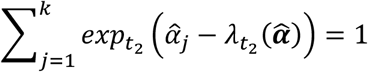.

Center loss is an optional setting in our model. Center loss was first proposed to assist models to learn discriminative representations in supervised learning.[19] Optimizing models with center loss is equal to minimizing intra-class variation defined by Eq. 6.

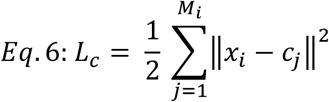

Where *c*_*i*_ is the centroid of *i*^*th*^cluster, and *x*_*j*_is the hidden features of *j*^*th*^sample in this cluster.

Because lowering center loss will push samples closer to the cluster center, the learned representations will be more discriminative in the hidden space. Though we did not use center loss to train models for Slide-seq data, we found that adding center loss during training can substantially improve accuracy in Fashion image data (Fig. S11) and the synthetic data with variance as 0.6. If a user has a biological dataset with some degree of known ground truth for comparison, initial data exploration should explore whether combining center loss and bi-tempered logistic loss is more appropriate to capture the known spatial features of the data.

#### 5. Normalized Mutual Information

Unlike supervised learning, we do not have ground truth for training in the CoSTA approach. To monitor how well training proceeds, we use normalized mutual information (NMI) to compare clustering labels before and after weight updating across training epochs. Increase of NMI during training indicates a decreased changing of clustering labels and thus suggests convergence of model. We cannot hold aside a validation set during CoSTA training. Therefore, NMI also serves as a metric of overfitting. Once we do not observe a large jump of NMI in consecutive epochs, we consider that the model has converged.

### 6. Experiments with Common Image datasets

While experimenting with MNIST handwritten, USPS-digit, and Fashion image datasets that come with true labels, we noticed that the CoSTA approach can learn to predict more true labels than the model that is just initialized and exceeds UMAP+GMM with pixel values as features (Fig. S11). For the Fashion image dataset, CoSTA was greatly improved after we add center loss with bi-tempered logistic loss as a whole loss function. However, the learning ability of CoSTA with these datasets is less than with supervised learning approaches (typically >95% accuracy). The highest accuracy we got is 0.961 (MNIST handwritten), 0.931 (USPS-digit) and 0.686 (Fashion), as measured by NMI between the clustering label and true class label. NMIs achieved with CoSTA applied to the MNIST and Fashion datasets are higher than for all other deep learning clustering methods, and the CoSTA NMI for USPS scores second in the ranking of deep learning approach performance.[7] We also tested whether SpatialDE can identify patterns in these three image datasets. We used the automatic histology pattern detection implemented in SpatialDE to cluster images in MNIST handwritten, USPS-digit, and Fashion into 10 groups, and SpatialDE achieved 0.532 (MNIST handwritten), 0.658 (USPS-digit), and 0.568 (Fashion) NMIs, which are even lower than UMAP+GMM clustering with pixels (Fig. S11).

### Simulation datasets

#### 1. Simulation of synthetic data with 5 patterns

We followed the simulation approach in SPARK to generate 10,000 fake genes that can be assigned into 5 distinct patterns.[5] We added residual errors onto each spatial coordinate independently based on a normal distribution with mean of zero and variance ranging from 0.2 to 0.6, resulting in 5 synthetic datasets with different noise levels. The simulation code can be found at https://github.com/xzhoulab/SPARK.

#### 2. Synthetic data with mask

We selected synthetic data with variance as 0.4 for this test. We arbitrarily selected a region to mask, a region in the blue circle of Fig. S3b. Though it is an arbitrary selection, we intentionally avoid any region that is crucial for each expression pattern. Therefore, the mask region will not disrupt spatial patterns visually. For each pattern, we randomly chose half of the simulated genes and added the mask by suppressing expression in that region to zero. The other half of the simulated genes remain intact. Thus, we have 5,000 genes each with and without masks.

#### 3. Mimic real spatial transcriptomic data by mixing SE and non-SE genes

We still focus on synthetic data with variance of 0.4. We further introduced non-SE genes to build more synthetic datasets that have different ratios of SE and non-SE genes. Code to generate non-SE genes can also be found at https://github.com/xzhoulab/SPARK. In this test, we generated 5 synthetic datasets with SE and non-SE ratios ranging from 90:10 to 10:90.

#### SE Gene Calling

To call out SE genes, we use an approach of ensemble learning. Simply put, we train 5 CoSTA models independently. We then calculate a set of nearest neighbors for every gene in the same dataset, using neighbor set sizes of 5, 10, 15, 20, 25, 30, 40, 50, and 100. This is because different genes with their neighbors may form a community with different sizes. Using a broad range of neighboring set sizes can enable us to include SE genes that only form a small community with a few genes as well as SE genes that fall into a large gene group. Next, we calculated Jaccard similarities across the 5 ConvNets and keep genes that have averaged Jaccard similarity larger than 0.2 at least in one level of neighbor set sizes: 5, 10, 15, 20, 25, 30, 40, 50, or 100.

### Correlated Gene Calling

To find significant correlated genes, we use the learned features from one of 5 CoSTA models to calculate Euclidean distance pairwise between all genes. For example, to get significant correlated genes with *Vim*, we calculated distances of all other SE genes to *Vim* based on the learned features. Then, we used these distances to create a null distribution. Distances that have Z-scores lower than -2.323 (p<0.01) are considered significant, and genes that have significant distances would be called out as correlated genes to *Vim*. Because we trained 5 independent models, we obtain 5 sets of correlated genes for each SE gene in the data. Then, we keep correlated genes that show up in at least in 3 models.

### MERFISH Analysis

We obtained the MERFISH dataset collected on the mouse preoptic region of the hypothalamus from Dryad[14](https://datadryad.org/stash/dataset/doi:10.5061/dryad.8t8s248), and we used the slice at Bregma + 0.11 mm from animal 18 for analysis as used for SPARK analysis.[5] We reduced the image resolution 10-fold and resized images to 85×85 matrices. Next, we directly applied a customized CoSTA model to the MERFISH dataset. This customized approach has the same general architecture that defines CoSTA, as described above. The customized ConvNet also has three convolutional layers but each convolutional layer has a larger filter, to reduce the overall size of the output. To compare with results from SPARK, we created null distributions for correlated gene calling by permuting images 100 times. Permuted images are forwarded through CoSTA to get permuted spatial features. Then we calculated their Euclidean distances with the spatial features of the true image, and these distances serve as the null distribution. Because the 9 defined cell type expression patterns are known, significantly correlated genes to these 9 expression patterns were called SE genes. For each gene in this MERFISH dataset, including the 5 blank controls, we calculated its Euclidean distances and its 100-time shuffled distances to the 9 expression patterns. If the true Euclidean distance of one gene to one cell type pattern are lower than Z-score -2.323, we call this gene an SE gene that is correlated to the expression pattern typical of this particular cell type. To visualize the training process, we project the feature vectors of each gene onto the first two UMAP dimensions and label each gene according to clusters defined using the whole feature vector. We use a linear intrinsic dimensionality (LID) estimator to quantify the change in cluster distinctness before and after training. This estimator mainly measures a ratio between distance of each datapoint to its the second closest datapoint and distance to its closest datapoint. Ratios are ordered from low to high and it fits a line that crosses the origin. The slope of this line represents the LID of this data in the latent space. Simply put, the lower LID, the more clustered datapoints are in the latent space. Indeed, among 10 different runs, spatial representations after training show lower LIDs than without training.

#### Analysis of Slide-seq with SPARK and SpatialDE

Analysis of Slide-seq with SPARK and SpatialDE follows the standard analysis pipelines proposed by these two methods, with default parameters. Code of analysis can be found at the GitHub repository (https://github.com/rpmccordlab/CoSTA).

## Figure Legends

**Supplementary Fig. 1**

Comparison of CoSTA and overlap analysis performance in finding correlated digits to digit 3. 1000 images are sampled from the full MNIST dataset, and each digit contains 100 samples. CoSTA (red bars) uniquely calls samples of digit 3 as correlated to digit 3. However, overlap analysis (blue bars) identifies some instances of all digits as showing some overlap with digit 3. CoSTA is more specific, but less sensitive: CoSTA reports a smaller number of correlated digit 3 images while overlap analysis reports a greater number of correlated digits overall.

**Supplementary Fig. 2**

Learning curve of CoSTA in true and shuffled synthetic datasets, with different variances. Clustering label generated by CoSTA is against true class label for measurement of NMI.

**Supplementary Fig. 3**

Performance of CoSTA using synthetic datasets with perturbations. a, Applying trained CoSTA to half and fully shuffled synthetic data. Genes are visualized by using spatial representation in 2D UMAP. Genes are colored based on the true synthetic pattern from which they are derived. Silhouette scores quantify how well the representation distinguishes different patterns. (Closer to 1 = more distinguishable patterns are recovered). b, Disruption test through masking. The masked region is circled in blue in the upper panel. Representation of pixelwise values (left) and features extracted by CoSTA (right) are visualized in 2D UMAP, and genes are colored based on pattern type from which they were generated (upper panel) or according to whether they belonged to the masked or unmasked set (lower panel).

**Supplementary Fig. 4**

Training CoSTA with different ratios of SE and non-SE genes. Genes are colored based on pattern membership (left) or SE type (right). Silhouette scores quantify how well the representation distinguishes different patterns, in SE and non-SE genes, respectively.

**Supplementary Fig. 5**

The number of overlapped neighbors of *Vim, Ctsd*, and *Gfap* before and after each weight updating across all epochs, considering either 10 nearest neighbors (left), 20 nearest neighbors (center), or 50 nearest neighbors (right).

**Supplementary Fig. 6**

The number of SE genes after 3 days and 2 weeks brain injury. a, Overlap of SE genes identified by SPARK, CoSTA learned with true data. b, learning curve of CoSTA with true and shuffled data. Y-axis shows NMI calculated between cluster labels at training epoch *t* and cluster labels at previous epoch *t-1*. X-axis shows training epoch *t*. c, Percent of all measured genes that are called SE genes by the 3 approaches.

**Supplementary Fig. 7**

Expression patterns of *Vim, Ctsd*, and *Gfap* 3 days and 2 weeks after brain injury. a, Expression patterns of *Vim, Ctsd*, and *Gfap* after 3 days after brain injury. b, Expression patterns of *Vim, Ctsd, Gfap* and genes involved in oligodendrocyte development (bottom row) 2 weeks after brain injury. Patterns that are visibly similar between *Vim, Gfap*, and *Ctsd* (small red boxes) are not strikingly visible in oligodendrocyte development genes.

**Supplementary Fig. 8**

Comparison of SPARK, SpatialDE, CoSTA, and pixel overlap results. a, Overlap of gene lists correlated with *Vim, Ctsd*, and *Gfap* at 2 weeks identified by CoSTA, SPARK, SpatialDE, and overlap analysis (“Slide-seq”). b, Examples of gene expression images for genes detected as similar to Vim, Gfap, and Ctsd by SPARK and also by CoSTA (left) or SPARK and not CoSTA (right). See Figure S7 for expression patterns of the query genes. All images are scaled between 0 and 1 for visualization purposes. Key visible regions of high expression in Vim, Gfap, and Ctsd are circled in red for cross comparison of all images. c, Gene Ontology term enrichment by CoSTA-ranked gene list (see Table S4) using GOrilla. d, Examples of gene expression images for genes highly ranked by CoSTA only (left), pixel only (middle), and both (right) as similar to Vim, Gfap, and Ctsd. Annotations next to gene names indicate rankings in CoSTA “C” and Pixel “P”.

**Supplementary Fig. 9**

Expression patterns of SE genes identified by CoSTA 2 weeks after brain injury. SE genes were clustered into 6 groups by SpatialDE and CoSTA. CoSTA cluster numbers correspond to Figure 4d and the most similar SpatialDE cluster is placed below the most closely corresponding CoSTA cluster when possible. The SpatialDE cluster containing *Vim, Gfap, and Ctsd* is cluster 2. Average expression pattern in 3^rd^ row shows the overall pattern of all genes combined in the 2-week dataset.

**Supplementary Fig. 10**

Effect of cluster number on CoSTA results with 2-week post injury Slide-seq data. a, SE genes identified by CoSTA with 10-100 clusters. b, CoSTA learning curve with 10-100 clusters. Y-axis shows NMI calculated between cluster labels at training epoch *t* and cluster labels at previous epoch *t-1*. X-axis shows training epoch *t*. c, Mean expression pattern of genes found to be correlated with *Vim, Gfap* and *Ctsd* identified by CoSTA with cluster numbers ranging from 10-

100. Raw count values are scaled from 0 to 1 for these visualizations.

**Supplementary Fig. 11**

CoSTA approach applied to clustering USPS, MNIST and Fashion datasets. Left panels: Models were trained for 10 epochs. After each weight updating, we clustered images into 10 clusters and directly compared them to true class labels through NMI. The grey line indicates clustering by UMAP+GMM with pixel values as features. The black line indicates clustering by SpatialDE. The orange line represents learning with combined center loss and bi-tempered logistic loss in Fashion dataset. Right panels: NMIs between clustering at the *t*^*th*^updating and the previous (*t* − 1)^*th*^updating.

**Supplementary Table 1**

Comparison of CoSTA and SpatialDE on 5 true and shuffled synthetic datasets. Adjusted Rand Index and Normalized Mutual Information are used to measure the ability of separating different spatial patterns. For shuffled data, each gene matrix still keeps its ground truth label but the original spatial pattern is disrupted.

**Supplementary Table 2**

Clusters of SE genes identified by CoSTA in the MERFISH dataset (cell type patterns are included in clusters).

**Supplementary Table 3**

Confusion matrix of clustering labels derived from CoSTA results compared to the original known experimental label.

**Supplementary Table 4**

All CoSTA SE genes ranked according to their similarity to query genes *Vim, Ctsd*, and *Gfap*. Ranking according to CoSTA feature vectors as well as by pixel correlation are shown. Columns 4 and 5 indicate whether each gene was found to be correlated to these query genes by SPARK or SpatialDE.

**Supplementary Table 5**

Genes in each cluster of SE genes detected by CoSTA from the 2-week Slide-seq data, as shown in figure 4b.

**Supplementary Table 6**

Runtime of CoSTA for 3-day and 2-week Slide-seq data. Runtimes are measured in minutes and under different numbers of clusters being assigned during training.

### Abbreviations

ConvNet: convolutional neural network
SE or SV gene: spatial expression or spatial variable gene
CoSTA: unsupervised ConvNet learning strategy for spatial transcriptomics analysis

## Declarations

### Ethics approval and consent to participate

Not applicable

### Consent for publication

Not applicable

### Availability of data and materials

The processed Slide-seq datasets were retrieved from https://singlecell.broadinstitute.org/single_cell/study/SCP354/slide-seq-study. We also deposited processed MERFISH and Slide-seq data and scripts for all analyses in this study at the GitHub repository (https://github.com/rpmccordlab/CoSTA) under an Open Source Initiative compliant MIT license. The version of the code used in the manuscript is available at DOI: 10.5281/zenodo.3948711.

## Author Contributions

Y.X. conceived the project, developed the computational approach, and performed all analysis. R.P.M. advised the project, and Y.X. and R.P.M. wrote the manuscript.

## Acknowledgments

We thank Tian Hong, Tongye Shen, and Amir Sadovnik for insightful discussion.

