## Supplementary Figures and Tables for "CoSTA: Unsupervised Convolutional Neural Network Learning for Spatial Transcriptomics Analysis"

Fig. S1

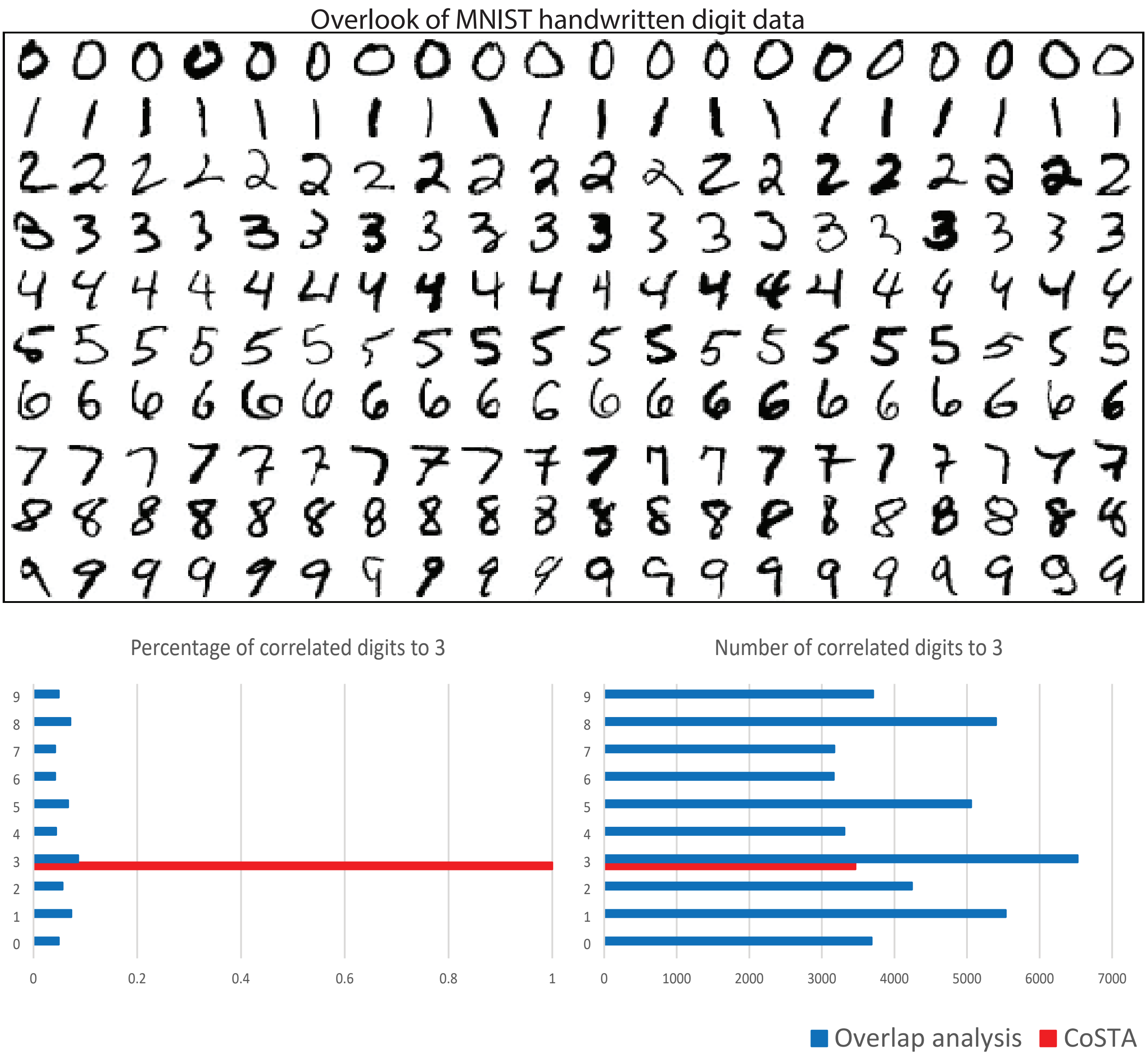
Overlook of MNIST handwritten digit data
Percentage of correlated digits to 3
Number of correlated digits to 3
9
8
7
6
5
4
3
2
1
0
0
0.2
0.4
0.6
0.8
1
0
1000
2000
3000
4000
5000
6000
7000
Overlap analysis
CoSTA

Fig. S2

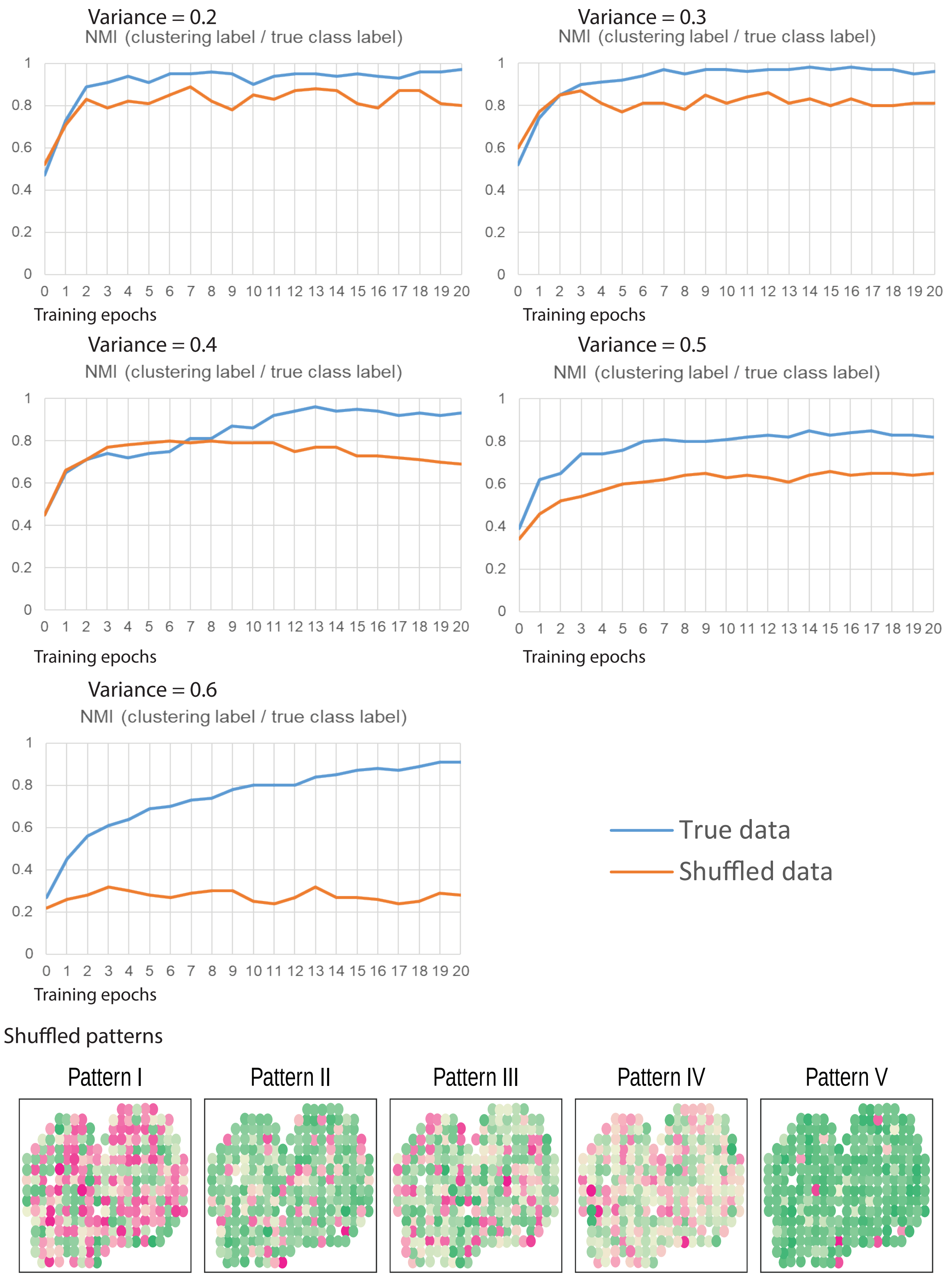
Variance = 0.2
NMI (clustering label / true class label)
Training epochs
Variance = 0.3
NMI (clustering label / true class label)
Training epochs
Variance = 0.4
NMI (clustering label / true class label)
Training epochs
Variance = 0.5
NMI (clustering label / true class label)
Training epochs
Variance = 0.6
NMI (clustering label / true class label)
Training epochs
True data
Shuffled data
Shuffled patterns
Pattern I
Pattern II
Pattern III
Pattern IV
Pattern V

Fig. S3

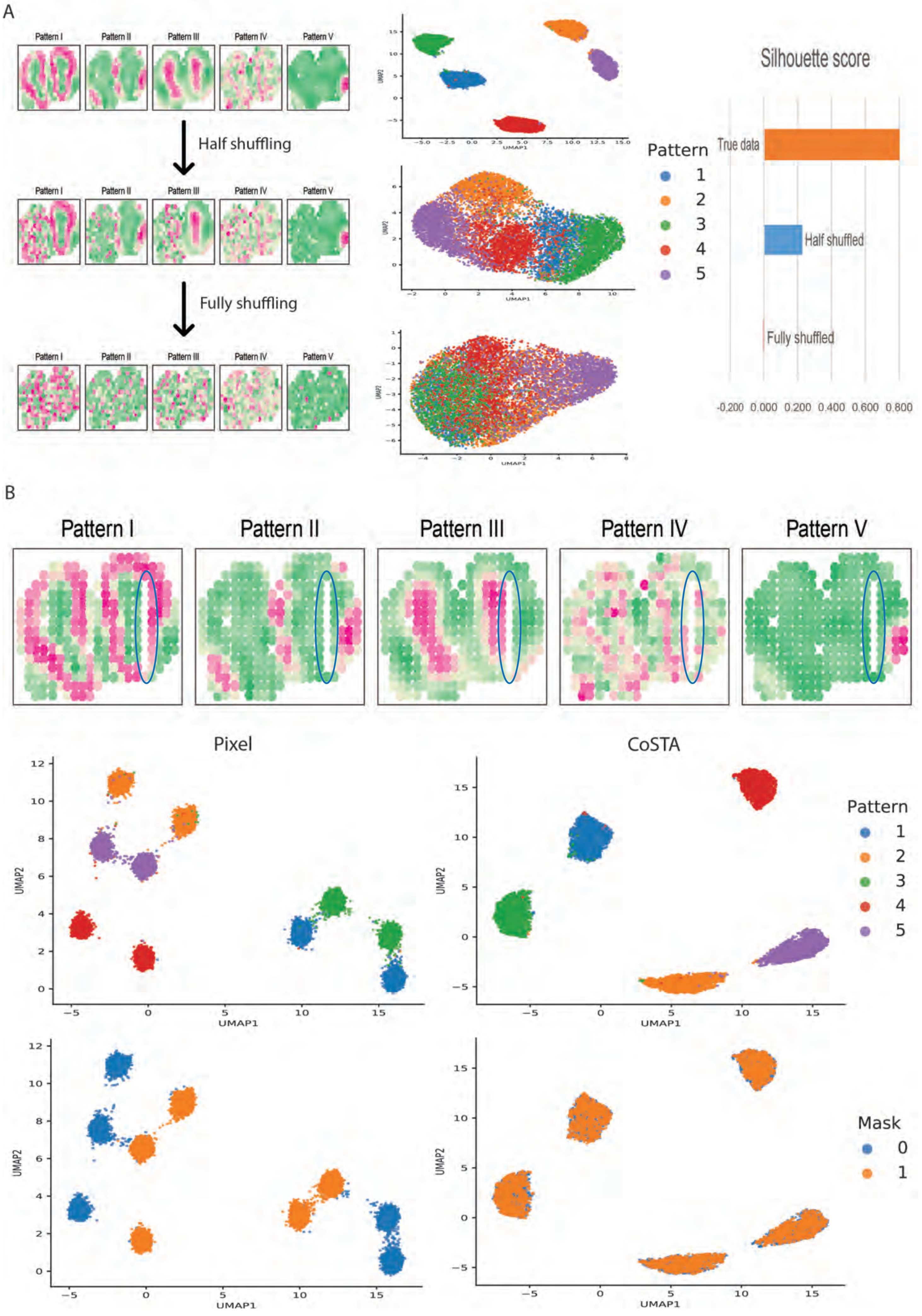
A
Pattern I
Pattern II
Pattern III
Pattern IV
Pattern V
Half shuffling
Fully shuffling
UMAP1
UMAP2
Pattern
1
2
3
4
5
Silhouette score
True data
Half shuffled
Fully shuffled
-0.200 0.000 0.200 0.400 0.600 0.800
B
Pattern I
Pattern II
Pattern III
Pattern IV
Pattern V
Pixel
CoSTA
Pattern
1
2
3
4
5
Mask
0
1
UMAP1
UMAP2

Fig. S4

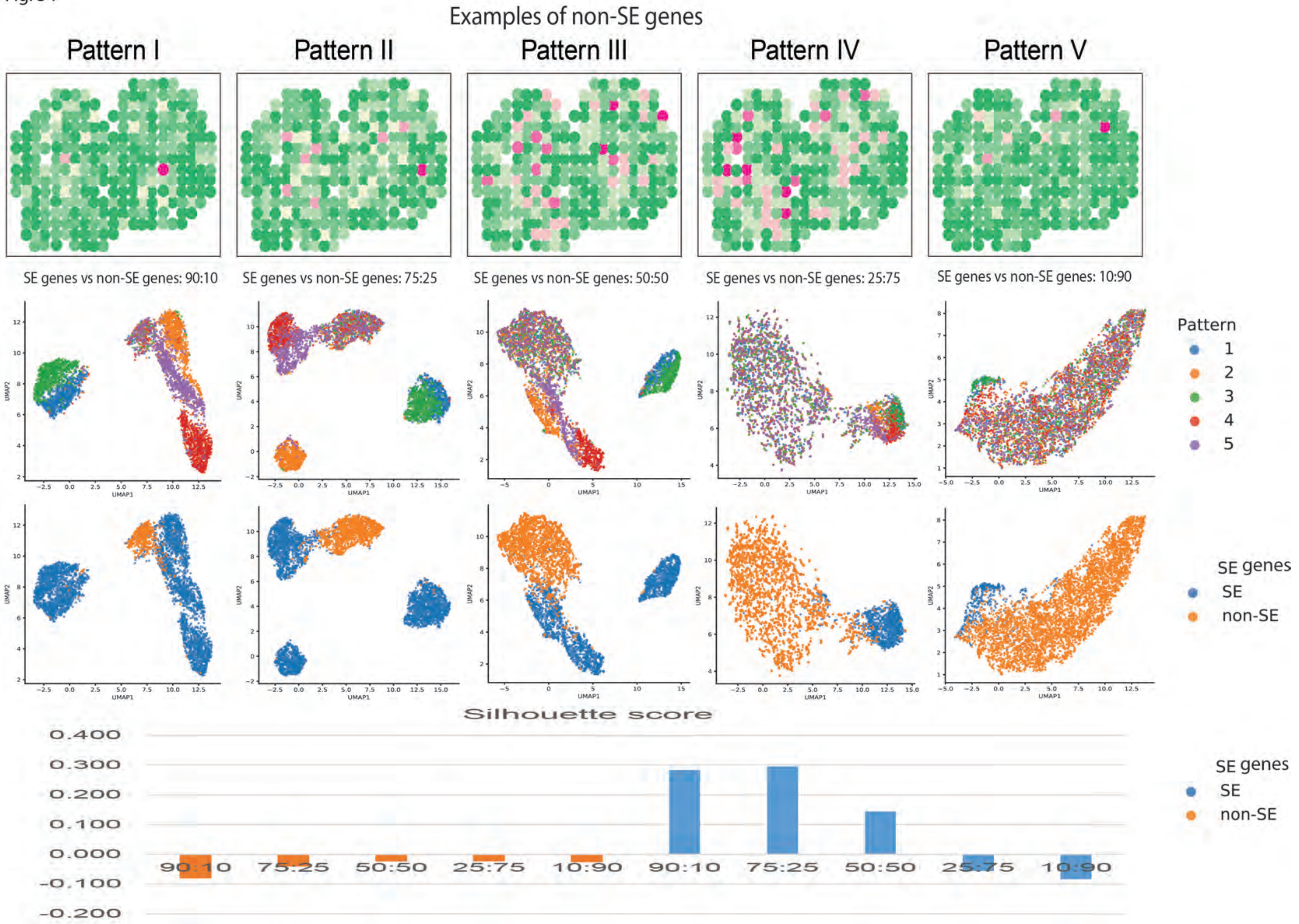
Examples of non-SE genes
Pattern I
Pattern II
Pattern III
Pattern IV
Pattern V
SE genes vs non-SE genes: 90:10
SE genes vs non-SE genes: 75:25
SE genes vs non-SE genes: 50:50
SE genes vs non-SE genes: 25:75
SE genes vs non-SE genes: 10:90
Pattern
1
2
3
4
5
SE genes
SE
non-SE
UMAP1
UMAP2
Silhouette score
0.400
0.300
0.200
0.100
0.000
-0.100
-0.200
90:10
75:25
50:50
25:75
10:90
90:10
75:25
50:50
25:75
10:90
SE genes
SE
non-SE

### Fig. S5

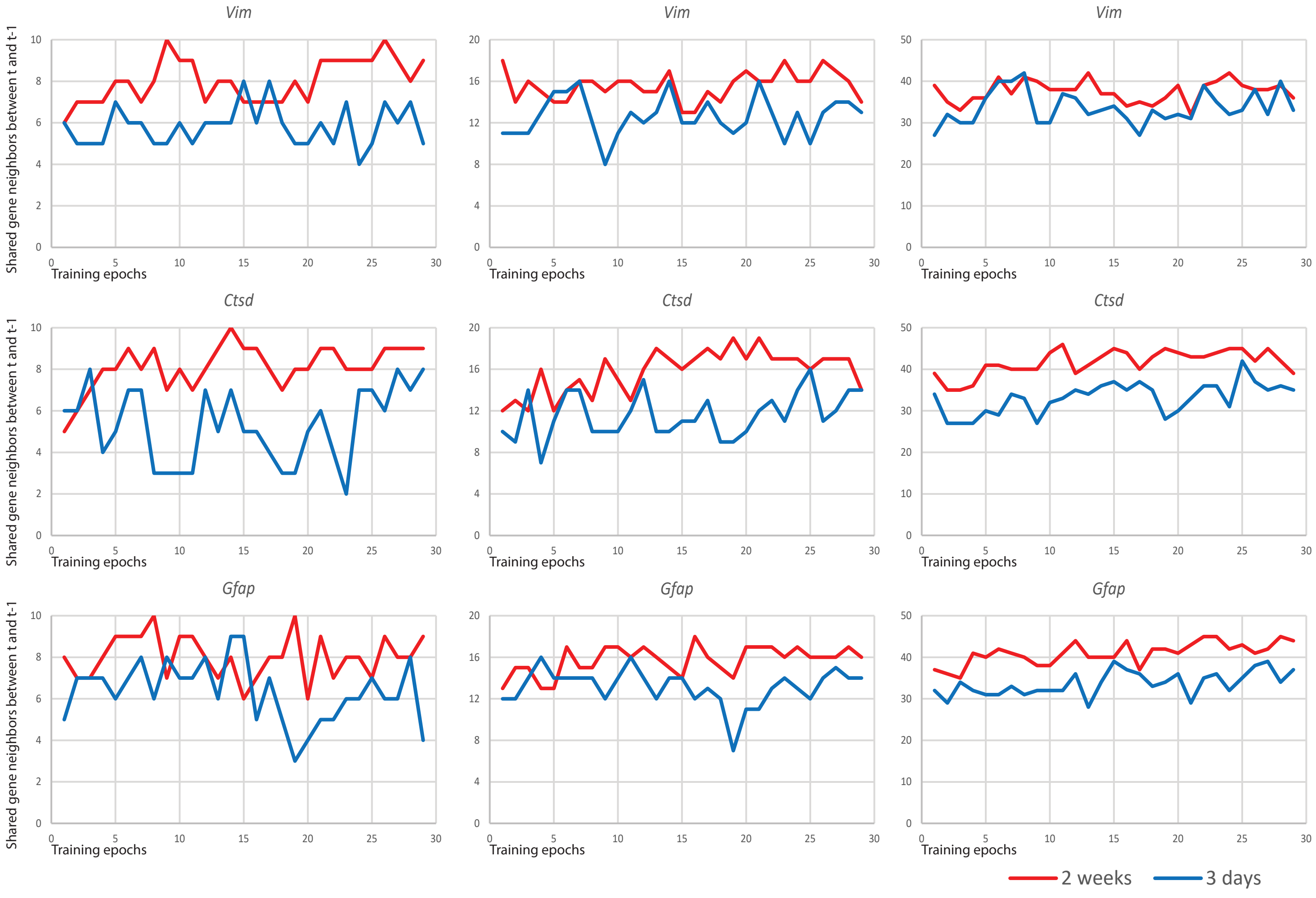
Vim
Ctsd
Gfap
Shared gene neighbors between t and t-1
Training epochs
2 weeks
3 days

Fig. S6

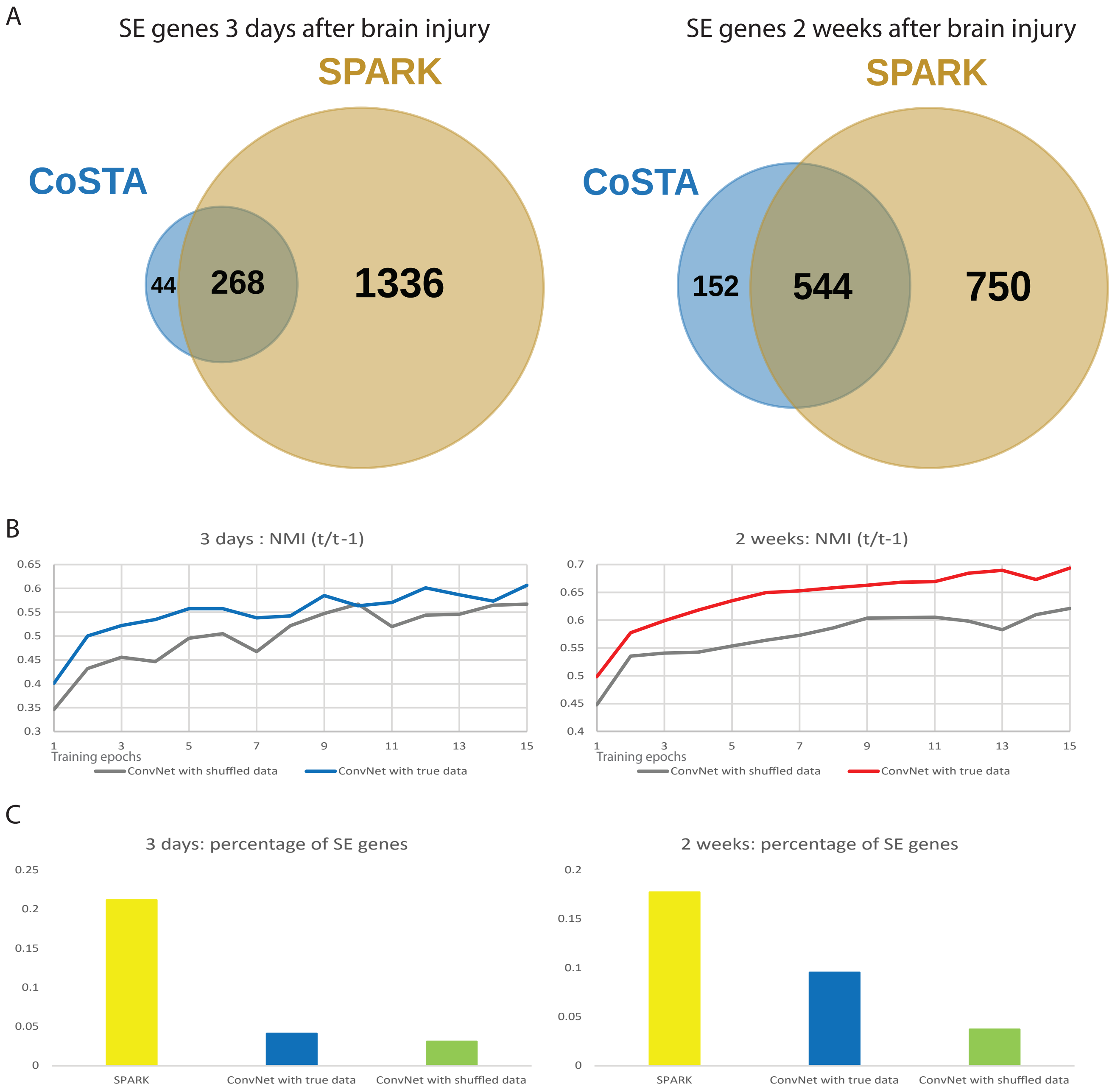
A
SE genes 3 days after brain injury
SPARK
CoSTA
44
268
1336
SE genes 2 weeks after brain injury
SPARK
CoSTA
152
544
750
B
3 days : NMI (t/t-1)
0.65
0.6
0.55
0.5
0.45
0.4
0.35
0.3
1 3 5 7 9 11 13 15
Training epochs
ConvNet with shuffled data
ConvNet with true data
2 weeks: NMI (t/t-1)
0.7
0.65
0.6
0.55
0.5
0.45
0.4
1 3 5 7 9 11 13 15
Training epochs
ConvNet with shuffled data
ConvNet with true data
C
3 days: percentage of SE genes
0.25
0.2
0.15
0.1
0.05
0
SPARK
ConvNet with true data
ConvNet with shuffled data
2 weeks: percentage of SE genes
0.2
0.15
0.1
0.05
0
SPARK
ConvNet with true data
ConvNet with shuffled data

Fig. S7

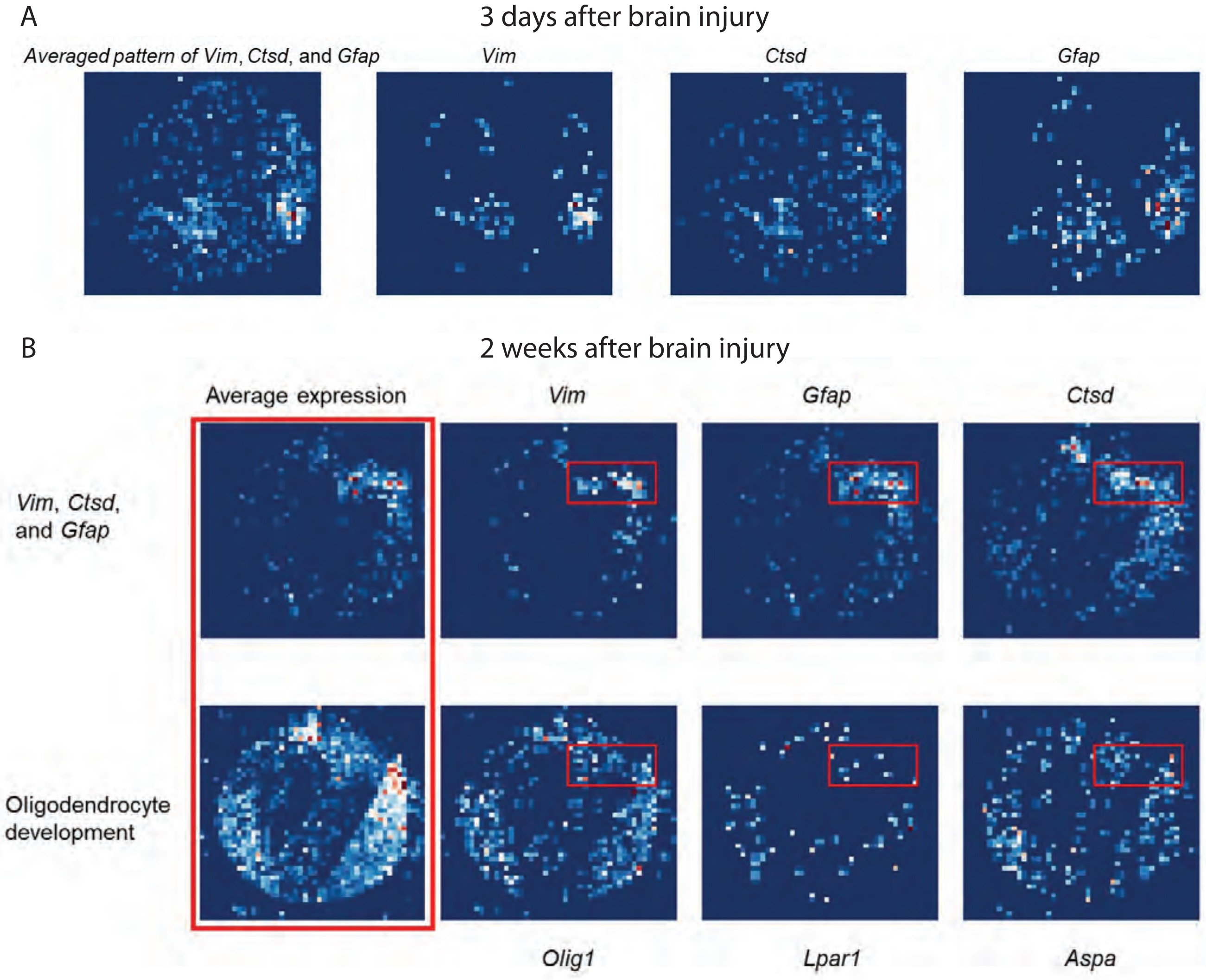
A
3 days after brain injury
Averaged pattern of Vim, Ctsd, and Gfap
Vim
Ctsd
Gfap
B
2 weeks after brain injury
Average expression
Vim
Gfap
Ctsd
Vim, Ctsd, and Gfap
Oligodendrocyte development
Olig1
Lpar1
Aspa

#### Fig. S8

**A**

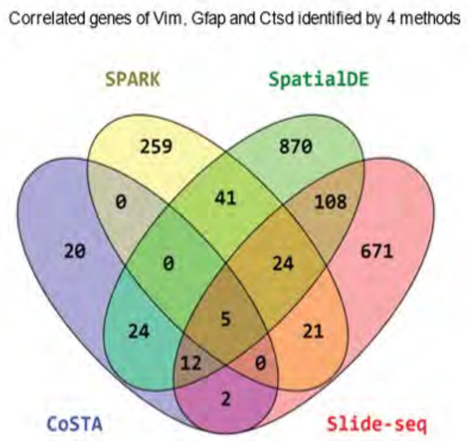


**B**

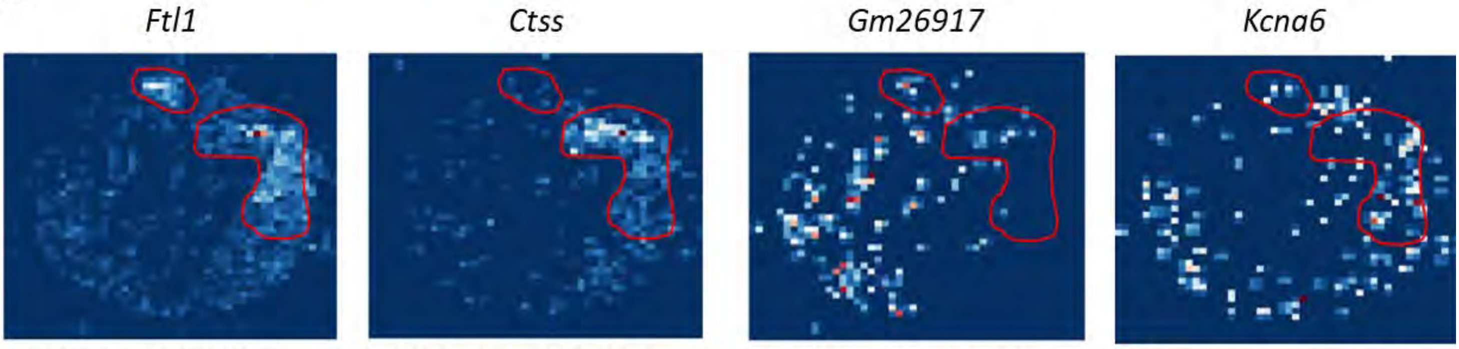


SPARK and CoSTA (#9), #313 by pixel

SPARK and CoSTA (#5), #134 by pixel

SPARK and NOT CoSTA (#578), #66 by pixel

SPARK and NOT CoSTA SE

**C**

| GO biological process | Fold Enrichment | FDR |
|---|---|---|
| collagen metabolic process | 89.70 | 0.008 |
| RNA metabolic process | 1.41 | 0.079 |
| Intermediate filament organization | 149.5 | 0.18 |
| Regulation of vascular endothelial growth factor signaling pathway | 74.75 | 0.26 |
| Astrocyte differentiation | 49.83 | 0.24 |

**D**

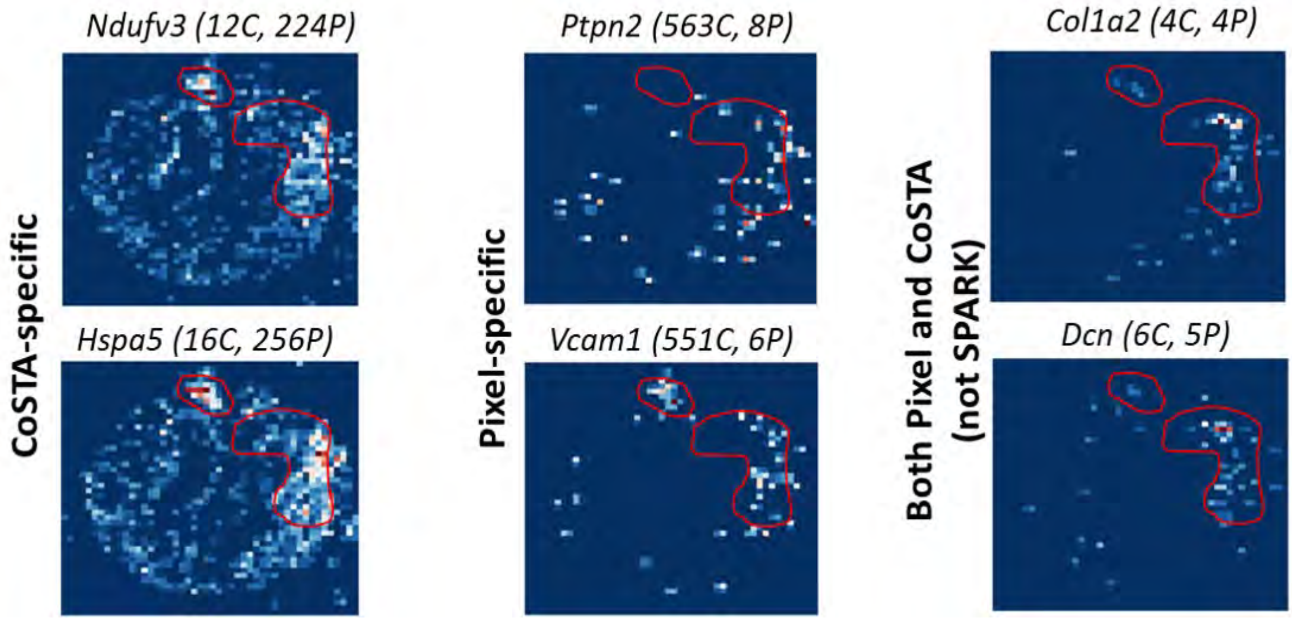

Fig. S9

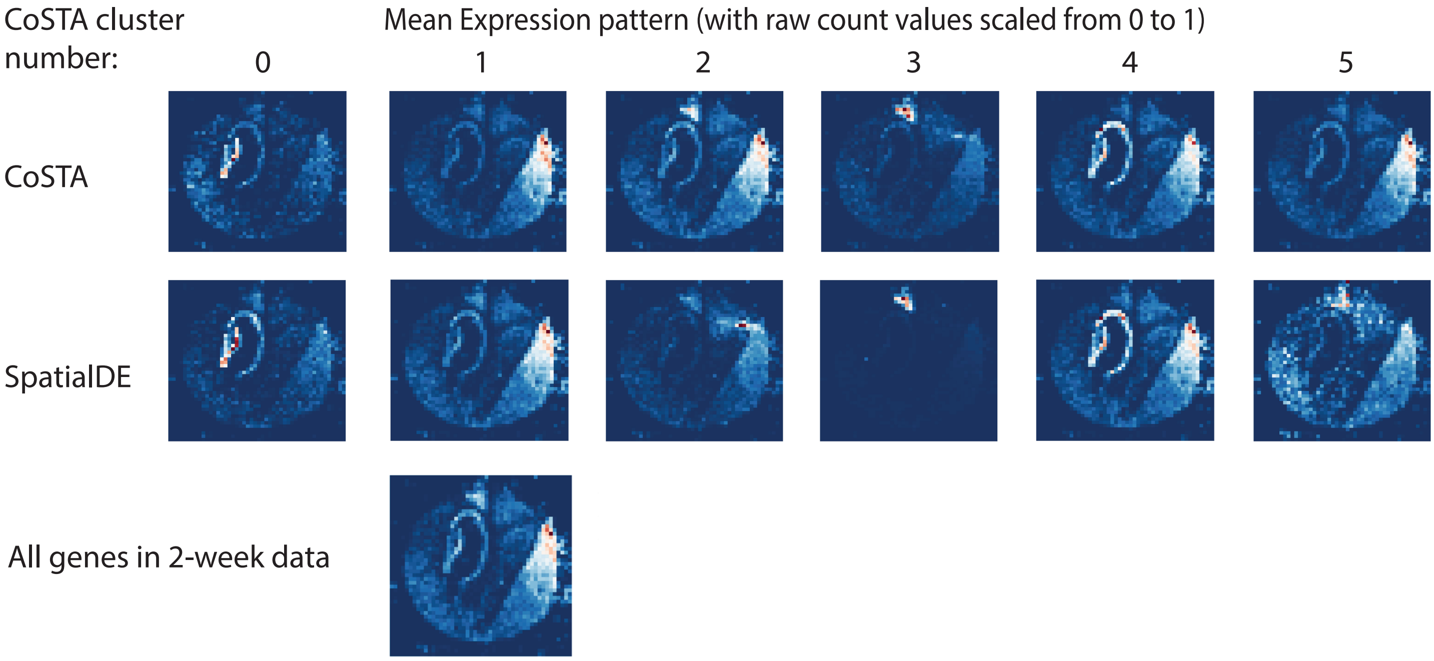
CoSTA cluster number:
Mean Expression pattern (with raw count values scaled from 0 to 1)
0
1
2
3
4
5
CoSTA
SpatialDE
All genes in 2-week data

Fig. S10

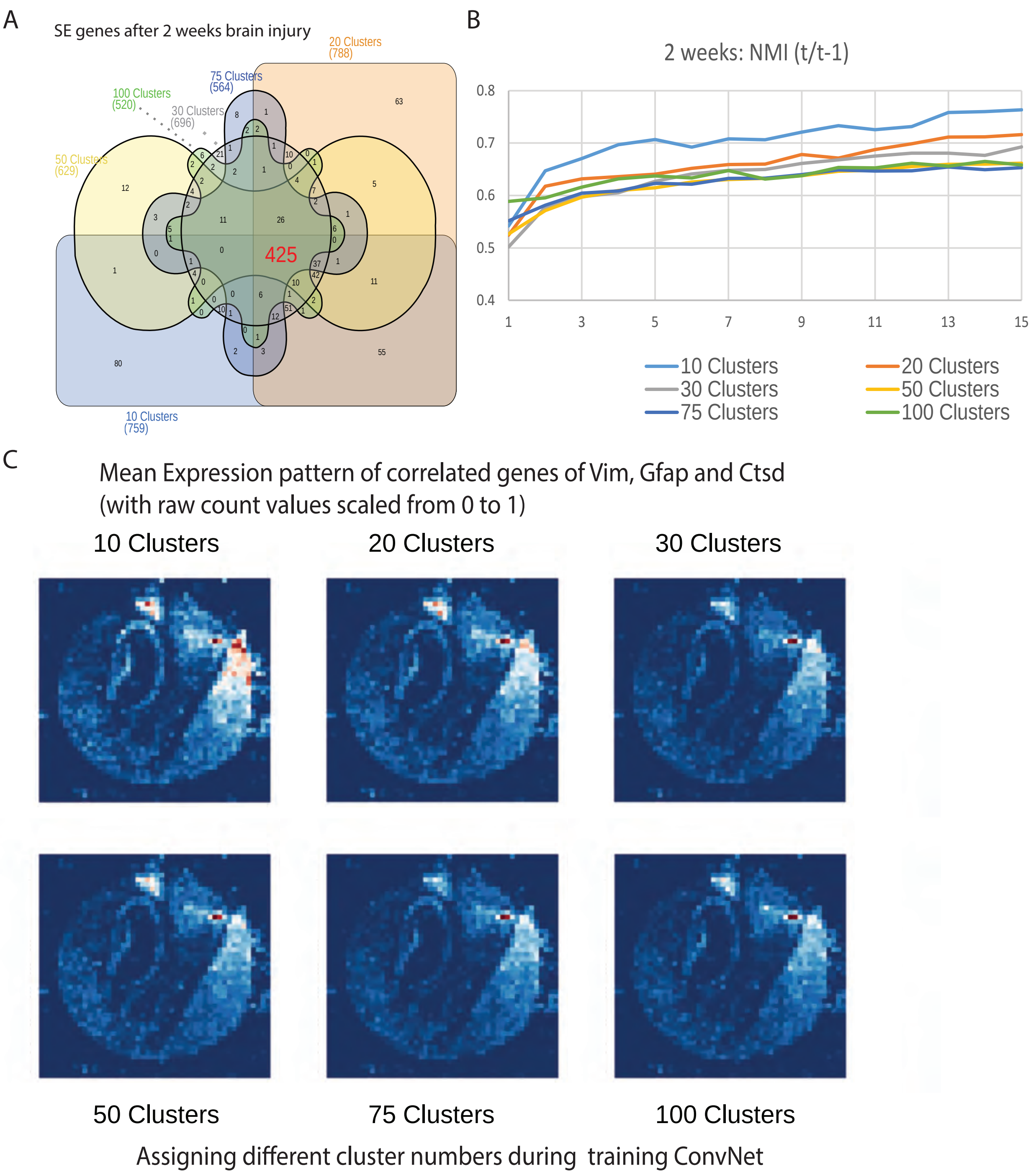

A
SE genes after 2 weeks brain injury
20 Clusters (788)
75 Clusters (564)
100 Clusters (520)
30 Clusters (696)
50 Clusters (629)
10 Clusters (759)
425
B
2 weeks: NMI (t/t-1)
0.8
0.7
0.6
0.5
0.4
1
3
5
7
9
11
13
15
10 Clusters
20 Clusters
30 Clusters
50 Clusters
75 Clusters
100 Clusters
C
Mean Expression pattern of correlated genes of Vim, Gfap and Ctsd
(with raw count values scaled from 0 to 1)
10 Clusters
20 Clusters
30 Clusters
50 Clusters
75 Clusters
100 Clusters
Assigning different cluster numbers during training ConvNet

Fig. S11

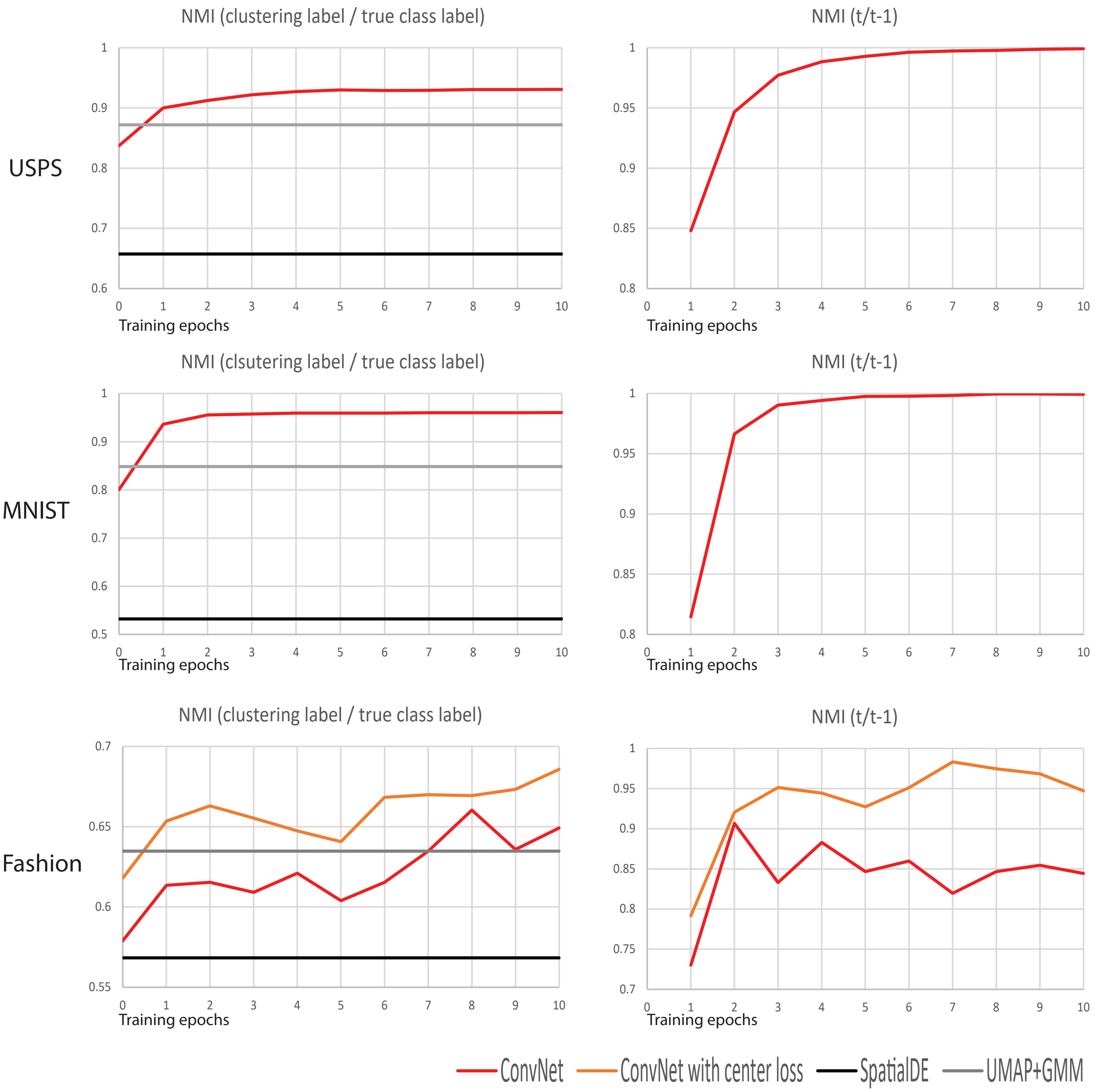

NMI (clustering label / true class label)
NMI (t/t-1)
USPS
NMI (clsutering label / true class label)
NMI (t/t-1)
MNIST
NMI (clustering label / true class label)
NMI (t/t-1)
Fashion
Training epochs
ConvNet
ConvNet with center loss
SpatialDE
UMAP+GMM

**Supplementary Table 1**

| NMI | CoSTA | | SpatialDE | |
|---|---|---|---|---|
| noise level (variance) | True data | Shuffled data | True data | Shuffled data |
| 0.2 | 0.97 | 0.89 | 1 | 0.86 |
| 0.3 | 0.98 | 0.87 | 0.99 | 0.99 |
| 0.4 | 0.96 | 0.8 | 0.99 | 0.99 |
| 0.5 | 0.85 | 0.66 | 0.98 | 0.98 |
| 0.6 | 0.52\|0.91 | 0.32 | 0.97 | 0.97 |

**Supplementary Table 2**

| Gene | Cluster |
|---|---|
| Endothelial 1 | 0 |
| Ermn | 0 |
| Gabra1 | 0 |
| Gjc3 | 0 |
| Igf1r | 0 |
| Man1a | 0 |
| Ndrg1 | 0 |
| OD Mature 2 | 0 |
| Sema3c | 0 |
| Sgk1 | 0 |
| Slco1a4 | 0 |
| Ttyh2 | 0 |
| Aldh1l1 | 1 |
| Amigo2 | 1 |
| Ar | 1 |
| Arhgap36 | 1 |
| Astrocyte | 1 |
| Cbln1 | 1 |
| Cbln2 | 1 |
| Cckar | 1 |
| Cpne5 | 1 |
| Creb3l1 | 1 |
| Crhr2 | 1 |
| Cspg5 | 1 |
| Dgkk | 1 |
| Excitatory | 1 |
| Gabrg1 | 1 |
| Galr1 | 1 |
| Glra3 | 1 |
| Gpr165 | 1 |
| Htr2c | 1 |
| Igf2r | 1 |
| Inhibitory | 1 |
| Irs4 | 1 |
| Isl1 | 1 |
| Kiss1r | 1 |
| Onecut2 | 1 |
| Oprd1 | 1 |
| Oprk1 | 1 |
| Oprl1 | 1 |
| Pak3 | 1 |
| Pnoc | 1 |
| Prlr | 1 |
| Rnd3 | 1 |
| Scg2 | 1 |
| Slc17a6 | 1 |
| Sox4 | 1 |
| Sox6 | 1 |
| Sox8 | 1 |
| Syt4 | 1 |
| Tmem108 | 1 |
| Adora2a | 2 |
| Bdnf | 2 |
| Brs3 | 2 |
| Ccnd2 | 2 |
| Chat | 2 |
| Endothelia | 2 |
| Gbx2 | 2 |
| Gem | 2 |
| Grpr | 2 |
| Krt90 | 2 |
| Lpar1 | 2 |
| Microglia | 2 |
| Nts | 2 |
| OD Mature | 2 |
| Rgs5 | 2 |
| Rxfp1 | 2 |
| Selplg | 2 |
| Cdkn1a | 3 |
| Cenpe | 3 |
| Cplx3 | 3 |
| Cyp19a1 | 3 |
| Fezf1 | 3 |
| Fn1 | 3 |
| Klf4 | 3 |
| Mbp | 3 |
| Ndnf | 3 |
| Necab1 | 3 |
| Ntng1 | 3 |
| Nup62cl | 3 |
| OD Mature | 3 |
| Opalin | 3 |
| Plin3 | 3 |
| Ramp3 | 3 |
| Slc17a8 | 3 |
| Sp9 | 3 |
| Sytl4 | 3 |
| Tacr1 | 3 |
| Calcr | 4 |
| Cxcl14 | 4 |
| Esr1 | 4 |
| Gad1 | 4 |
| Gal | 4 |
| Gda | 4 |
| Npy2r | 4 |
| Penk | 4 |
| Rgs2 | 4 |
| Serpinb1b | 4 |
| Th | 4 |
| Trhr | 4 |
| Coch | 5 |
| OD Immat | 5 |
| Pcdh11x | 5 |
| Pdgfra | 5 |
| Traf4 | 5 |
| Crhbp | 6 |
| Cyr61 | 6 |
| Ebf3 | 6 |
| Endothelia | 6 |
| Fst | 6 |
| Gnrh1 | 6 |
| Lmod1 | 6 |
| Mki67 | 6 |
| Myh11 | 6 |
| OD Immat | 6 |
| OD Mature | 6 |
| Oxt | 6 |
| Pericytes | 6 |
| Sst | 6 |
| Syt2 | 6 |
| Tac2 | 6 |
| Ucn3 | 6 |
| Adcyap1 | 7 |
| Aqp4 | 7 |
| Avpr1a | 7 |
| Cckbr | 7 |
| Cd24a | 7 |
| Ependyma | 7 |
| Etv1 | 7 |
| Fos | 7 |
| Mlc1 | 7 |
| Nnat | 7 |
| Nos1 | 7 |
| Npy1r | 7 |
| Omp | 7 |
| Pou3f2 | 7 |
| Sema4d | 7 |
| Slc15a3 | 7 |
| Slc18a2 | 7 |
| Sln | 7 |
| Tac1 | 7 |
| Tiparp | 7 |
| Avpr2 | 8 |
| Egr2 | 8 |
| Galr2 | 8 |
| Pgr | 8 |
| Synpr | 8 |
| Vgf | 8 |

**Supplementary Table 3**

| | **0** | **1** | **2** | **3** |
|---|---|---|---|---|
| **0** | 2266 | 36 | 2 | 6 |
| **1** | 1 | 5117 | 115 | 157 |
| **2** | 0 | 78 | 7396 | 102 |
| **3** | 4 | 114 | 91 | 7085 |

Clustering label

Experiment label

**Supplementary Table 5**

| Cluster 0 |
|---|
| Cdkn1b |
| Vps13c |
| Hap1 |
| Wbscr17 |
| Scn3b |
| Pitpnm2 |
| Gm26917 |
| Marcksl1 |
| Jph1 |
| Vav3 |
| Slc17a6 |
| C1ql2 |
| Sema5a |
| Nr3c2 |
| Prox1 |
| Rasl10a |
| Limd2 |
| Plk5 |
| Ahcyl2 |
| Epha7 |
| Tcf7l2 |
| Fam163b |
| Vamp1 |
| Dock10 |

| Cluster 1 | | |
|---|---|---|
| Gpm6b | Psma7 | Dlgap1 |
| Ssb | Prrc2c | Cbx5 |
| Lmo4 | Peg3 | Prkar1b |
| Prdx2 | Sltm | Scn1b |
| Rock2 | Ldha | Elavl4 |
| Gria3 | Soga3 | Prkacb |
| Cmip | Phactr1 | Atp2b2 |
| Ppp1r9a | Tuba4a | B2m |
| Rtf1 | Ankrd12 | Lpgat1 |
| Vamp2 | Srrm2 | Ndrg3 |
| Snap47 | Myo5a | Sncb |
| Arpp19 | Strbp | Dclk1 |
| Ensa | Klf9 | Gnb1 |
| Cfap36 | Egr1 | Trim37 |
| Ube2k | Ntrk2 | Sult4a1 |
| Pdap1 | Zfr | mt-Tp |
| Rsrp1 | Map1a | Fam81a |
| Tuba1b | Luc7l3 | Pak1 |
| Rabep1 | Kif21a | Kifap3 |
| Acot7 | Stmn1 | Gabra1 |
| Zc3h13 | R3hdm1 | Pde1a |
| Chd3os | Sptbn1 | Ddx24 |
| Syngr1 | Fam171b | |
| Mapt | Eef1a2 | |
| Eid1 | Atp6v0e2 | |
| Nrn1 | 6330403K07Rik | |
| Rgs4 | Ptprn | |
| Zranb2 | Mphosph8 | |
| Nars | 2210016L21Rik | |
| Zfand5 | Rbm25 | |
| Eif5b | Zfp365 | |
| Mapk10 | 3110035E14Rik | |
| Atp5o | Rangap1 | |
| Ntm | Ttyh1 | |
| Prkcb | S100b | |
| Zmynd11 | Sf3b1 | |
| Psmc1 | Plcb1 | |
| Basp1 | Pcmt1 | |
| Fam168a | Gda | |
| Oxct1 | Ncl | |
| Rufy3 | Rph3a | |
| Apba2 | Klc1 | |
| Ndufb9 | Mdh2 | |
| Cdk5r1 | Atp1a2 | |
| Ghitm | Srpk2 | |
| Ndufa10 | Tmem50a | |
| Ppig | Sars | |
| Atp5c1 | Nefm | |
| Aplp1 | Rrp1 | |
| Ndrg2 | Son | |
| Scd2 | Pgm2l1 | |
| Srsf11 | Nrxn2 | |
| Tagln3 | Cplx1 | |
| Sbno1 | Eif3c | |
| Uqcc2 | Ctsb | |

| Cluster 2 |
|---|
| Dnaja1 |
| Matr3 |
| Dnm1 |
| Snap25 |
| Rab6a |
| Cox4i1 |
| Arpp21 |
| Gria2 |
| Ywhag |

| Cluster 3 | |
|---|---|
| Trf | Car14 |
| Xpnpep3 | Pbxip1 |
| Ppp1r1b | Hemk1 |
| Sox5 | Igfbp7 |
| Gcnt2 | Ier3 |
| Tmem98 | Cldn11 |
| Cryab | C1qb |
| Phactr2 | Stk39 |
| Otx2 | Cd63 |
| Acaa2 | Ezr |
| Atox1 | Aldh2 |
| Ifi27 | Cox8b |
| Qdpr | C1qc |
| Haus2 | Rbp1 |
| Ccnd2 | Vat1l |
| Slc31a1 | Slc16a2 |
| Crhbp | Vim |
| Gpr88 | Gfap |
| Hist1h2bc | Tgfb2 |
| Cnp | Lyz2 |
| Itpk1 | Lars2 |
| Col1a2 | Spint2 |
| Igfbp2 | Ctnnal1 |
| Etfb | Npc2 |
| Tnfaip8 | Dcn |
| Ranbp9 | Trpm3 |
| Slco1c1 | Calb2 |
| Plin3 | Mag |
| Elovl7 | Cdkn1c |
| Krt12 | Gng5 |
| Ctsd | Sh3d19 |
| Pcp4l1 | Sostdc1 |
| Ctss | Vamp8 |
| Tesk1 | Pvrl3 |
| Kl | |
| Kcnj13 | |
| Mgp | |
| S100a1 | |
| Tbc1d9 | |
| Pltp | |
| Cd59a | |
| Calml4 | |
| Ccdc66 | |
| Cab39l | |
| Gas6 | |
| Nwd2 | |
| Clic6 | |
| Mal | |
| Nnat | |
| Igf2 | |
| Folr1 | |
| Fxyd1 | |
| 1500015O10Rik | |
| Slc16a6 | |
| Prlr | |

| Cluster 4 | |
|---|---|
| Cpne6 | Chgb |
| Grin2a | Cfl1 |
| Mycbp2 | Npm1 |
| Arpc1a | Nfib |
| Enc1 | Ywhaz |
| Xist | Ncdn |
| Wasf1 | Cacna1e |
| Gnaq | Ppp3r1 |
| Hpca | Atp1a3 |
| Zeb2 | Btbd9 |
| Epha4 | Cttnbp2 |
| Neurod6 | Ap2a2 |
| Olfm1 | Ppfia2 |
| Camk2b | Kif5c |
| Gria1 | Ndufb7 |
| Herc1 | Ppp3cb |
| Grin2b | Ank3 |
| Syn2 | Brinp1 |
| Auts2 | Camkk1 |
| Ubxn4 | Mapk1 |
| Nptx1 | Capza2 |
| Erc2 | Gabra5 |
| Thra | Nell2 |
| Zbtb20 | Bdnf |
| Kalrn | Nbea |
| Bcl11b | Napa |
| Cnih2 | Ywhah |
| Celf2 | Dynll1 |
| Gng2 | Ncam1 |
| Camta1 | Abr |
| Sh3bgrl3 | Ptk2b |
| Rab2a | Rbfox1 |
| Lppr2 | Fam131a |
| Arf3 | Rnf112 |
| Syne1 | Neurod2 |
| Epha5 | Nptxr |
| Cpne4 | Ak5 |
| Trim2 | Snca |
| Ddx5 | Cadm2 |
| Synj1 | |
| Sepw1 | |
| Ogfrl1 | |
| Pnmal2 | |
| Zbtb18 | |
| Nrxn1 | |
| Tubb2a | |
| Rap1gds1 | |
| 2010300C02Rik | |
| Wipf3 | |
| Arpc5 | |
| Stxbp6 | |
| Schip1 | |
| Sirt3 | |
| Pfn1 | |
| Mrfap1 | |

| Cluster 5 | | |
|---|---|---|
| Dlgap4 | Itpr1 | Smap1 |
| Atl1 | Hlf | Elavl3 |
| Dgkz | Hpcal4 | Igfbp6 |
| Ktn1 | Nlk | Prpf40b |
| Prkcz | Phactr3 | Nudt4 |
| Ndfip2 | Tbl1x | Ppap2b |
| Rora | Flywch1 | Tsnax |
| Ccl27a | Fam107a | Fgf12 |
| Cdc5l | Oxr1 | Luc7l |
| Fnbp1l | Rapgef4 | Ube2r2 |
| Plcb4 | Btbd10 | Serpine2 |
| Pik3r1 | Emc4 | Lamp5 |
| Pmm1 | Slc1a3 | Fabp3 |
| Chga | Nos1ap | Kif3a |
| Cdk11b | Car10 | Zfp91 |
| D430041D | Pcdh7 | Pfkm |
| Sv2b | Satb1 | Ndufaf2 |
| Ccdc186 | Clstn3 | Camk2g |
| Arhgap32 | Srp72 | Ldb2 |
| Phf3 | Stx1a | |
| Psmd2 | Akap8l | |
| Mgll | Rabl6 | |
| Pin1 | Nap1l2 | |
| Meis2 | Necap1 | |
| Rims2 | Rufy2 | |
| Srrm3 | Slc39a10 | |
| Golga4 | Nktr | |
| Snw1 | Kcnb1 | |
| Nemf | Pcdh9 | |
| Thoc2 | Gabra3 | |
| Clasp2 | Chd5 | |
| Rims1 | Kcna2 | |
| R3hdm2 | Gpbp1 | |
| Tubb4b | Bend6 | |
| Htatsf1 | Sgtb | |
| Ttc9b | Scrn1 | |
| Glrx2 | Cfdp1 | |
| Cabp1 | Homer1 | |
| Nr3c1 | Cobl | |
| Ddx1 | Lingo1 | |
| Stau2 | Asap1 | |
| Map9 | Pak1ip1 | |
| Foxp1 | Dnajc21 | |
| Wdr26 | 1700025G04Rik | |
| Sept11 | A830010M20Rik | |
| Pacsin1 | Add1 | |
| Scn1a | Lin7a | |
| Cacng2 | Tia1 | |
| Ankrd11 | Usp7 | |
| Cxxc5 | Gm10419 | |
| Zfp148 | Esf1 | |
| Cep290 | Epb4.1l3 | |
| Smarcc1 | Apc | |
| Rap2a | Camkk2 | |
| Pip5k1c | Rcan2 | |

**Supplementary Table 6**

| **Slide-seq** | **3-day** | **2-week** |
|---|---|---|
| # of genes | 7576 | 7294 |
| size of image | 48X48 | 48X48 |
| runtime (in min) | 11.5 | 12.5 |

(running with 30 clusters)

(running with assigning different clusters)

| 2-week | 10 clusters | 20 clusters | 30 clusters | 50 clusters | 75 clusters | 100 clusters |
|---|---|---|---|---|---|---|
| runtime (in min) | 8 | 9.5 | 12.5 | 14.5 | 21 | 29.5 |

| | |
|---|---|
| CPU | Intel i9-9880H |
| Memory | 64GB |
| GPU | NVIDIA Quadro T2000 |
